# Diverging repeatomes in holoparasitic Hydnoraceae uncover a playground of genome evolution

**DOI:** 10.1101/2024.12.09.626742

**Authors:** Woorin Kim, Nicola Schmidt, Matthias Jost, Elijah Mbandi Mkala, Sylke Winkler, Guang-Wan Hu, Tony Heitkam, Stefan Wanke

## Abstract

- The nuclear genomes of parasitic plants have undergone unique evolutionary trajectories to adapt to the heterotrophic lifestyle. These adaptations often involve large genomic alterations, potentially driven by repetitive DNAs. Despite the recognized role of repetitive DNAs for shaping plant genomes, their contribution to parasitic plant genome evolution remains largely unexplored. This study presents the first analysis of repetitive DNAs in eleven genomes of Hydnoraceae, a holoparasitic plant family.
- The repetitive DNAs were identified and annotated *de novo*. Patterns of abundance and presence-absence were aligned with the phylogenetic relationships, geographical distribution, and host shifts, suggesting repetitive DNAs’ key role in shaping Hydnoraceae genomes.
- The two Hydnoraceae genera, *Hydnora* and *Prosopanche*, exhibit distinct profiles. Eight *Hydnora* genomes are largely populated by long terminal repeat retrotransposons, while three *Prosopanche* genomes vary greatly in repetitive DNA composition, including *P. bonacinae* with massive amplifications of a single DNA transposon and *P. panguanensis* with over 15% 5S rDNA, compared to <0.1% in some Hydnoraceae. Both extremely low and very high abundance of 5S rDNA challenges our current understanding on chromosomal stabilization and rDNA transcription.
- These genome dynamics, while inviting for further investigations, suggest rapidly evolving repeat profiles, potentially accelerated by the adaptation to the parasitic lifestyle.

## Introduction

Parasitic plants, representing approximately 1% of angiosperms, have taken an evolutionary turn from autotrophy to heterotrophy which entailed considerable genomic alterations (Nickrent, 2020; Lyko & Wicke, 2021; Chen *et al*., 2023). Holoparasitic plants, which completely lack photosynthetic activity, possess a highly reduced plastid genome (plastome), due to the gene losses of photosynthetic and other functional genes like the plastid-encoded tRNAs (Lyko & Wicke, 2021; Kim *et al*., 2023). Mitochondrial genomes in parasitic plants show diverse evolutionary patterns across parasitic lineages, such as extensive gene losses in hemiparasitic mistletoes (*Viscum*) (Zervas *et al*., 2019), maintenance of typical gene set in some holoparasitic dodders (*Cuscuta*) compared to autotrophic angiosperms (Petersen *et al*., 2020), as well as high structural complexity alongside many integrations of foreign sequences in the holoparasitic Balanophoraceae (Zhou *et al*., 2023). Although nuclear genomic studies are available for only a limited number of parasitic lineages, first findings suggest deeply intertwined host-parasite relation (Davis & Wurdack, 2004; Yang *et al*., 2019), accelerated evolutionary rates (Lemaire *et al*., 2011; Bromham *et al*., 2013), and massive convergent gene loss with concomitant gene enrichment associated with the parasitic invasion (Sun *et al*., 2018; Xu *et al*., 2022; Ashapkin *et al*., 2023; Chen *et al*., 2023). Furthermore, changes in genome size and chromosomal structures in parasitic plants suggest a potential role of repetitive DNAs in shaping their genomes (Neumann *et al*., 2021).

Repetitive DNAs (repeats) are a major component of all plant nuclear genomes (Orozco-Arias *et al*., 2019). Repeats significantly contribute to genome dynamics by driving mutations in coding sequences, influencing chromosomal organization, and affecting genome size (Raskina *et al*., 2008; Mehrotra & Goyal, 2014; Lee & Kim, 2014; Anderson *et al*., 2019). However, regarding parasitic plants genomes, studies on repeats has been limited to a few taxa, such as *Orobanche* and *Striga* (Orobanchaceae), *Cuscuta* (Convolvulaceae), and Rafflesiaceae (Weiss-Schneeweiss *et al*., 2006; Piednoël *et al*., 2015; Neumann *et al*., 2021; Cai *et al*., 2021). These studies uncovered long terminal repeat retrotransposons (LTR retrotransposons) of the Ty3-*gypsy* and Ty1-*copia* types as the most abundant repeats in aforementioned parasitic lineages, mirroring patterns seen in autotrophic plants (Piednoël *et al*., 2015; Neumann *et al*., 2021; Cai *et al*., 2021). The accumulation of repeats, e.g. retrotransposons or satellite DNAs, is often linked to genome size expansions (Neumann *et al*., 2021; Schmidt *et al*., 2024). In *Cuscuta* genomes, the loss of a specific class of Ty3-*gypsy* LTR retrotransposons may be associated with the formation of holocentric chromosomes (Neumann *et al*., 2021). Given the potential impact of repeats on genome dynamics, further investigation on the repetitive fraction of the genome (repeatome) within and across parasitic plant lineages is essential to investigate their role in parasitic plant genome evolution (Lyko & Wicke, 2021).

The focus naturally extends to Hydnoraceae, a family of root holoparasites within the Magnoliid order Piperales (Nickrent *et al*., 2002; Jost *et al*., 2021), whose nuclear genomes still remain to be explored. Hydnoraceae comprise the two genera *Hydnora* (Thunberg, 1775) and *Prosopanche* (Bary, 1868), which are distributed across the Old and New World, respectively. *Hydnora* species are found from Southern Africa to the Arabian Peninsula, while *Prosopanche* species are found throughout South and Central America (Gómez P. *et al*., 1981; Musselman & Visser, 1989; De Carvalho *et al*., 2021; Hatt *et al*., 2024b). The most common host plant families of Hydnoraceae are Fabaceae and Euphorbiaceae, beside up to 16 other families that have been reported as hosts (Hatt *et al*., 2024a).

Recent taxonomic studies recognize ten species of *Hydnora* and seven species of *Prosopanche*, with a more finely resolved classification dividing the genus *Hydnora* into four subgenera (Hatt *et al*., 2023, 2024b). Their unusual lifestyle as root holoparasites, spending most of their lifetime underground (Tennakoon *et al*., 2007), and extremely reduced morphology suggest distinctive evolutionary dynamics in their genomes. Although molecular genomic studies on Hydnoraceae are scarce, existing research suggests an accelerated evolutionary rate in the nuclear-encoded 18S ribosomal DNA (Nickrent & Duff, 1996; Lemaire *et al*., 2011; Bromham *et al*., 2013). The plastid genomes of Hydnoraceae are highly reduced, and exhibit extremely long branches in a phylogenetic tree, even when compared to other parasitic plants (Naumann *et al*., 2013; Jost *et al*., 2020). On the other hand, the mitochondrial genome of *Hydnora visseri* is four-fold larger than those of related species (Yu *et al*., 2023).

Here, we aim to illuminate the genome evolution of holoparasitic Hydnoraceae, focusing on the repetitive DNAs. To do so, we analyse low coverage Illumina sequences of eight *Hydnora* and three *Prosopanche* species, which cover about 80% and 40% of the species diversity, respectively. We use read clustering to deduce the individual repeat profiles of the eleven Hydnoracea, using *H. visseri* as a reference species. The additional long reads of *H. visseri* allowed us to reconstruct representative sequences of the most abundant repetitive DNAs and to provide detailed repeat classifications, serving as a basis for a comparative repeatome analysis of Hydnoraceae. Furthermore, the ribosomal DNAs from Hydnoraceae genomes were analyzed in-depth to elucidate the sequence variation and genomic organization, as well as variable genomic abundance within and between both Hydnoraceae genera.

## Material and Methods

### Plant material, DNA extraction and genome sequencing

Eleven Hydnoraceae taxa were sequenced after extracting DNA from silica dried plant material as described by (Naumann *et al*., 2013; Jost *et al*., 2020, 2022; Mkala *et al*., 2023). In short, DNAs were extracted using the CTAB method (Doyle & Doyle, 1987), modified to include an RNase A (Thermo Scientific, Waltham, MA, USA) treatment (10 mg/ml), except for *H. visseri*, *H. longicollis,* and *P. americana*, where the DNeasy Plant Maxi Kit (Qiagen, Venlo, Netherlands) was used instead. Library preparation was conducted using NEBNext Ultra DNA kits for the most samples, except for *H. abyssinica* and *P. americana*, where the Illumina TruSeqDNA kit was used. Sequencing was performed to generate paired-end reads using the following Illumina platforms: HiSeq-2000 (for *H. visseri* and *H. abyssinica*), HiSeq Rapid Mode (for *H. longicollis*, *H. africana, H. triceps, H. hanningtonii, H. solmsiana, H. esculenta, P. bonacinae, P. panguanensis*), and NextSeq High Output (for *P. americana*). For *H. visseri,* long read*s* were generated from DNA extracted using the CTAB method described above. One continuous long-read (CLR) single molecule real-time (SMRT) Pacific Biosciences library was prepared from high-molecular-weight genomic DNA, following the ‘20 kb template preparation using Blue Pippin^TM^ size selection system’ protocol (version P/N 100-286-000-07). The final large-insert library was selected for fragments larger than 10 kb using the BluePippin^TM^ device (Sage Science Inc., Beverly, MA, US). This PacBio library was sequenced on 43 PacBio RS2 SMRT cells v3 for 6 hours using DNA polymerase P6 and sequencing chemistry 4.0. A full sampling list including the origin of the material is presented in Table S1.

### Global repeat analysis by read clustering

To prepare the input data for read clustering analyses, the quality of each set of Illumina reads was first evaluated using FastQC v.0.11.5 (Andrews, 2010). Subsequent read pre-processing involved several steps, including filtering, trimming, and random subsampling: Plastome sequences and Illumina adapters were filtered using Bowtie2 v.2.2.6 (Langmead & Salzberg, 2012). From each forward- and reverse-read dataset, five million reads were randomly sampled and trimmed to 100 nucleotides (nt) using Trimmomatic v0.39 (Bolger *et al*., 2014). Finally, the reads were merged into paired-end read files, each labeled with species-specific codes (see Table S1). For a comparative purpose, a close photoautotrophic relative of the Hydnoraceae, *Aristolochia fimbriata* (Aristolochiaceae, SRR13748080) underwent the same procedure and was included in the analysis. For the visual overview of the processes, see Fig. S1A.

Read clustering analysis for each Hydnoraceae species was performed using the RepeatExplorer2 pipeline (RE2; Novák *et al*., 2013) applying default parameters (90% similarity over 55% of the read length, cluster size threshold = 0.01% of the analyzed reads, minimal cluster size for assembly = 5). The REXdb (Viridiplantae version 3.0) database served as a reference for identifying transposable element protein domains for primary annotations. In addition, a comparative analysis was performed using a concatenated dataset of 1 million randomly subsampled reads for each species from the datasets used for individual read clustering analysis. A custom repeat database containing the most abundant repeats of the *H. visseri* genome (see next paragraph) was included for the individual and comparative read clustering analyses.

### *In silico* reconstruction of reference retrotransposons and 35S rDNA

17 reference repeats were reconstructed for the 32 most abundant read clusters, exceeding 0.2% of the analyzed reads, in the *H. visseri* genome (Table S1). To achieve this, the highest read-depth contigs of the corresponding read clusters provided by the pipeline were used for a basic local alignment search (BLASTn embedded in Geneious v6.1.8; Kearse *et al*., 2012) against *H. visseri* PacBio long reads (Table S2). The searched 20 best-matching long reads, representing the top 20 BLASTn hits with the lowest e-values, were investigated using self- and pairwise dotplots based on the presence of conserved protein domains for transposable elements or ribosomal RNA genes. The 20 full or fragmented repeats were then reconstructed as a consensus using MUSCLE alignments (Edgar, 2004), providing a reference repeat. Lastly, the conserved protein domains harbored by the reference repeats were annotated using DANTE (Novák *et al*., 2024). For the visual overview of the processes, see Fig. S1B. These reconstructed repeats were collected to create a custom repeat database for the annotation of repeats during subsequent RepeatExplorer2 analyses.

Further species-specific repetitive elements were reconstructed or obtained, including an unclassified repeat found in the genomes of *H. hanningtonii*, *H. abyssinica* and *H. solmsiana.* The highest read-depth contigs from the most abundant unclassified read cluster of each species (provided by the pipeline) were refined by mapping the all available short reads of the respective species to each contig using Bowtie2, generating consensuses. These three consensuses were aligned using MUSCLE (for alignment, see Fig. S2). Two unclassified repeats in *H. esculenta* genome were reconstructed using the same method. In addition, a *P. panguanensis*-specific satellite DNA and two *P. bonacinae*-specific satellite DNA candidates were obtained as consensus sequences provided by the pipeline (for alignment, see Fig. S3). The reconstructed repeats as well as the consensuses provided by the pipeline are found in public data repository (see Data availability).

### *In silico* reconstruction of Hydnoraceae 5S ribosomal RNA genes

For only three of the investigated Hydnoraceae species (*H. triceps*, *P. bonacinea*, and *P. panguanensis*), consensus sequences for the 5S rDNA were provided by the pipeline. These RE2 consensuses were refined by mapping the short reads from each, *H. triceps*, *P. bonacinea*, and *P. panguanensis* against the RE2 consensuses using Bowtie2 v.2.2.6 (Langmead and Salzberg 2012). The mapped reads to each RE2 consensus yielded improved 5S rDNA reference for *H. triceps*, *P. bonacinea*, and *P. panguanensis,* respectively. These 5S rDNA references were used for the reconstruction of the 5S rDNA of the remaining Hydnoraceae species: Short reads of the *Hydnora* species were mapped individually against the 5S rDNA reference from *H. triceps*, whereas *P. americana* short reads were mapped against the 5S rDNA reference from *P. panguanensis* using Bowtie2 v.2.2.6 (the respective number and proportion of mapped reads is listed in Table S3).

To compare the reconstructed Hydnoraceae 5S rDNAs with those from 56 other angiosperms, we extracted the 5S rDNA gene from available genome assemblies or whole genome sequencing data (for the list of the data, see Table S4). For each angiosperm species, a 5S rDNA consensus was generated from the alignment of 50 5S rDNA sequences extracted using BLAST, using *A. fimbriata* 5S rDNA gene as a query sequence. This query sequence was reconstructed using *A. fimbriata* Nanopore long reads (accession number: SRR13748081), based on a MAFFT alignment (Katoh *et al*., 2002) of 50 different 5S rRNA genes identified from the *A. fimbriata* long reads. For some species, 5S rDNAs gene were retrieved from a publicly available rRNA database (Szymanski *et al*., 2002; http://combio.pl/rrna/). The sequence similarity of all 5S rDNAs to each other was visualized with a distance matrix generated using the Kimura-2-parameter model (Kimura, 1980).

Additionally, a complete 5S monomer (including the 5S rRNA gene and the intergenic spacer) was reconstructed for *H. visseri*. To do so, *H. visseri*-derived PacBio long reads were mapped to the *H. visseri* 5S rDNA (see above) using Bowtie2 v.2.2.6. Repetitive 5S monomers were identified on 37 long reads by inspecting self- and pairwise dotplots. In total, 248 partial and full-length monomers were extracted and aligned using MUSCLE to create the consensus sequence of the complete 5S rDNA monomer. The arrangement of the 5S rDNA monomers on the PacBio long reads was investigated using BLAST (Altschul *et al*., 1990; threshold E-value 1E-5). The BLAST hits on each long read were visualized with a zoom factor of 500 on each self-dotplot created by FlexiDot (word size 10; Seibt *et al*., 2018).

## Results

### The *H. visseri* genome is dominated by Ty3-*gypsy* retrotransposons of the Tekay and Ogre type

To gain insights into the genomic repeat profiles of the holoparasitic Hydnoraceae, the preliminary read clustering analysis of *H. visseri* was investigated thoroughly, ultimately providing a repeat database containing the most abundant repetitive elements from the *H. visseri* genome (Fig. 2). Using this database, Hydnoraceae repeat profiles were annotated in detail, allowing for an in-depth individual and comparative genomic analysis. For our reference taxon *H. visseri*, it is estimated that over half of its genome consists of repetitive sequences (56.3%; Fig. 2A). Ty3-*gypsy* retrotransposons are the most abundant repeats, accounting for two-thirds of the *H. visseri* repeatome, particularly Tekay and Ogre elements (Fig. 2A, B). Reference sequences were reconstructed for the 17 most abundant repeats, including *Hydnora*Tekay1-4 (T1, T2, T3, T4), *Hydnora*Ogre1-5 (O1, O2, O3, O4, O5), *Hydnora*CRM1 and 2 (C1, C2), *Hydnora*Angela1 and 2 (A1, A2), *Hydnora*SIRE1 and 2 (S1, S2), *Hydnora*Galadriel1 (G1), and a complete 35S rDNA monomer comprising the 18S, 5.8S, 26S genes alongside intergenic spacers (Table S1). Among these 17 repeats, 12 belong to the Ty3-*gypsy* retrotransposons (i.e. Tekay, Ogre, CRM and Galadriel). A few Tekay and Ogre elements predominate the repeatome (T1, O1, T2; Fig. 2B), and the reconstructed sequences of the two Tekay elements retain *gag* and *pol* open reading frames (Fig. 2C). On the other hand, Ty1-*copia* retrotransposons account for 2.4% of the *H. visseri* genome, represented mainly by Angela and SIRE (Fig. 2B; Table S2). The 35S rDNA contributes 1.4% to the genome, whereas 5S rDNA was not detected by the pipeline. These detailed annotations of *H. visseri* read clustering analysis was enabled using the reconstructed repeats, which provided a basis to further annotate other Hydnoraceae repeatomes.

**Figure 1.**
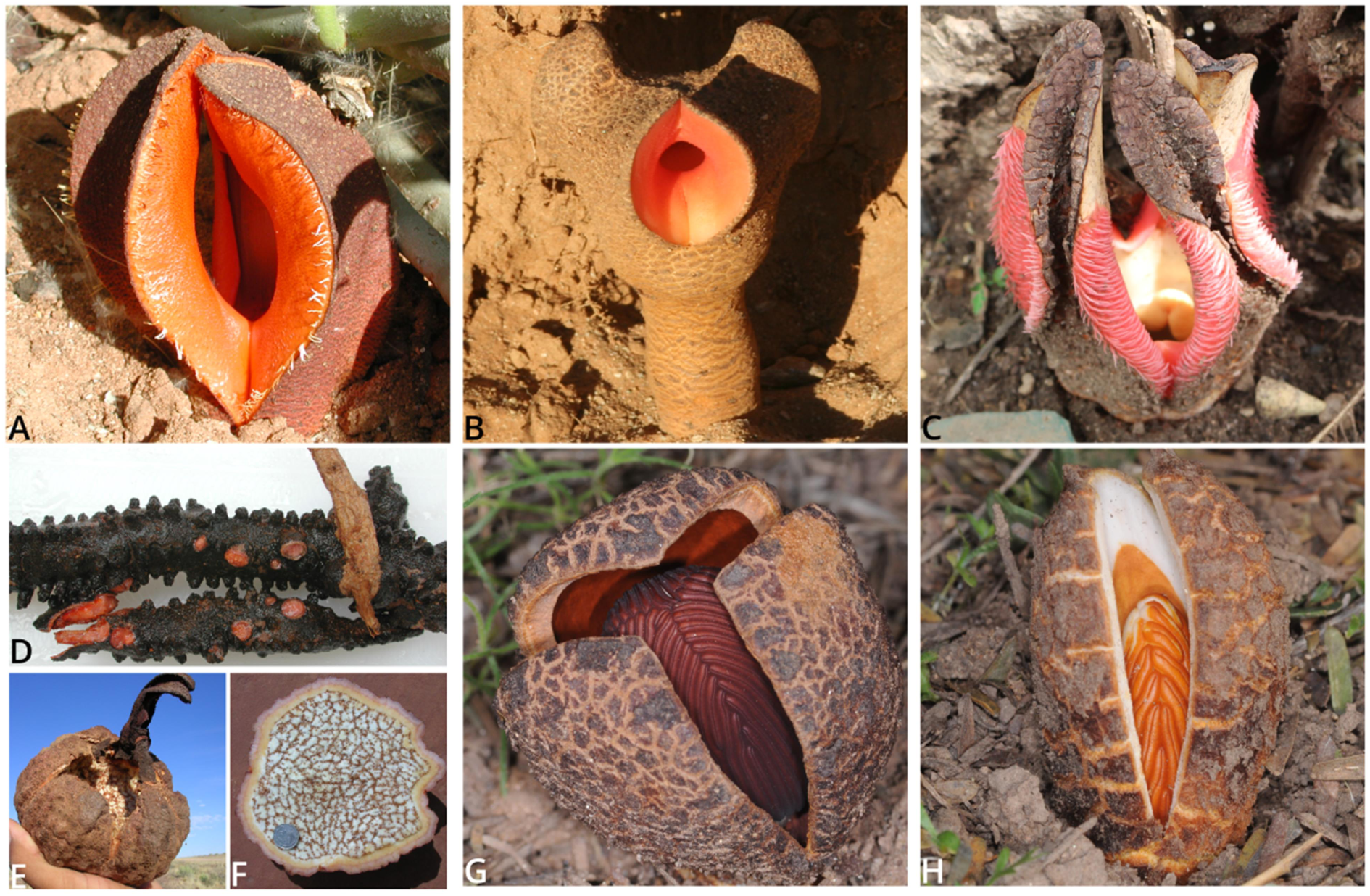
Flowers, fruit, and rhizome of Hydnoraceae; A. *H. visseri* flower; B. *H. triceps* flower; C. *H. abyssinica* flower; D. *H. visseri* rhizome; E. *H. visseri* fruit; F. *H. visseri* fruit section; G. *P. americana* flower; H. *P. bonacinae* flower; Photographs (A, D, E, F) by Jay Bolin, (B) by Stefan Wanke, (C) by Elijah Mbandi Mkala (G, H) by Andrea Coccuci.

**Figure 2.**
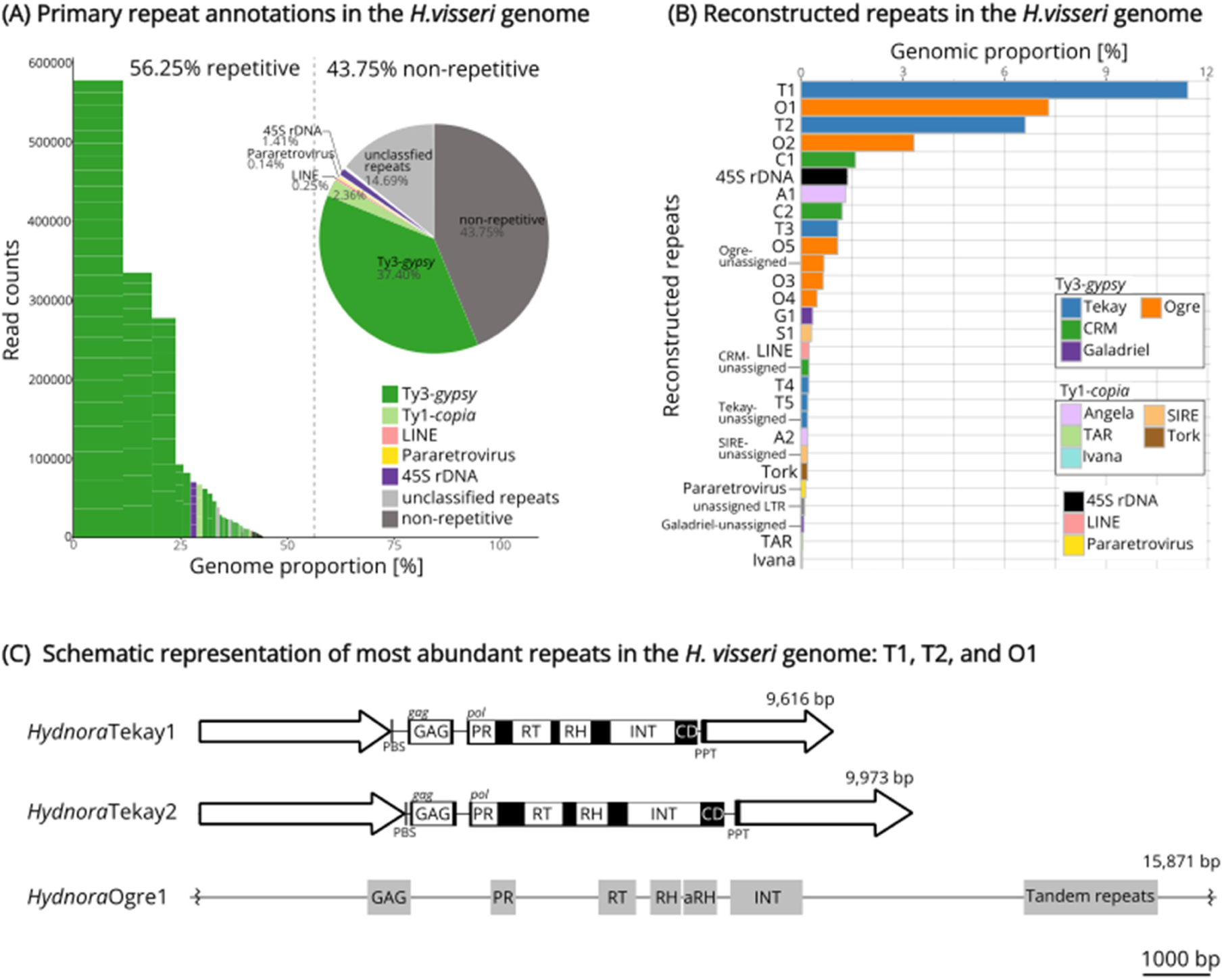
(A) The relative proportion of repetitive elements in the *H. visseri* genome, primarily annotated by the RepeatExplorer2 pipeline. Bars indicate superclusters, with each stack within the bar representing a read cluster. The height and width of the bars correspond to the read counts of the supercluster and the genomic proportion, respectively. The proportions are calculated based on the number of reads in each category divided by the total number of analyzed reads. (B) Genomic proportions of reconstructed repeats from the *H. visseri* genome. The bar length represents the genomic proportions, calculated as the percentage of reads within the corresponding clusters divided by the total number of analyzed reads. Abbreviations of the individual LTR retrotransposons are as follows: *Hydnora*Tekay1-5) T1-T5, *Hydnora*Ogre1-5) O1-O5, *Hydnora*CRM1-2) C1-C2, *Hydnora*Angela1-2) A1-A2, *Hydnora*Galadriel1) G1, *Hydnora*SIRE1) S1 (Table S2). Bars labeled as ‘unassigned’ and further categories (i.e. ‘Tork’, ‘Pararetrovirus’, ‘TAR’, and ‘Ivana’) correspond to the totality of elements belonging to the respective category and falling below the abundance threshold for the reconstruction of reference elements (proportions were added up). (C) Schematic representation of most abundant repeats in the *H. visseri* genome. Long Terminal Repeats (LTRs) are represented by arrows at the 5’ and 3’ ends of the elements. The protein domains were annotated using DANTE, using default setting (REXdb: Viridiplantae_v.3.0; Scoring Matrix: BLOSUM80). The shaded black blocks behind the protein annotations of *Hydnora*Tekay1 and *Hydnora*Tekay2 refer to the open reading frames (ORFs), where the *gag* and *pol* ORFs are separated. The zigzag lines on the 5’ and 3’ ends of *Hydnora*Ogre1 indicate truncated ends. PBS: primer binding site, PPT: polypurine tract, GAG: gag-like protein, PR: protease, RT: reverse transcriptase, RH: ribonuclease H, INT: integrase, CD: chromodomain, aRH: archeal ribonuclease H.

### Repeat compositions of genus *Hydnora* mirror that of *H. visseri*, whereas those of genus *Prosopanche* highly diverge

To capture the genome dynamics represented in repeat compositions, the individual repeatomes of the *Hydnora* and *Prosopanche* genomes were analyzed and annotated using the repeat database from the *H. visseri* read clustering analysis (Figs 3 and 4). The *Hydnora* genomes show variability in their overall repeat content, ranging from 34.6% in *H. esculenta* to 56.7% in *H. longicollis* (Fig. 3; Table S2). Ty3-*gypsy* retrotransposons are the most abundant repeats in all analyzed *Hydnora* genomes (9.3% to 40.2%; Table S5). In particular, different Ty3-*gypsy* elements predominate the genomes: Tekay elements are most abundant in the genomes of *H. visseri* (T1), *H. longicollis* (T1)*, H. triceps* (T1)*, H. africana* (T1), and *H. esculenta* (T2), while Ogre elements predominate in the genomes of *H. hanningtonii* (O1), *H. abyssinica* (O1), and *H. solmsiana* (Ogre; Fig. 4A-G). In addition, there are highly abundant unclassified repeats in the genomes of *H. hanningtonii*, *H. abyssinica*, and *H. solmsiana*, that turned out to be the same repeat present in all three genomes (indicated as arrows in Fig. 4D-F; for the alignment, see Fig. S2). The *H. esculenta* genome contains two additional, highly abundant unclassified repeats (Fig. 4G). Aside from the high abundance of Ty3-*gypsy* retrotransposons, *Hydnora* genomes are characterized by scarce amounts of 5S rDNA, DNA transposons, and satellite DNAs (Fig. 4A-G; Table S5). In contrast, these elements are much more abundant in the *Prosopanche* genomes (Fig. 4H-J). 5S ribosomal DNA is the most prevalent repeat in the *P. panguanensis* genome, followed by satellite DNA (Ppan_Sat01; Fig. 4I, Table S6). The En/Spm_CACTA superfamily of DNA transposons, which contains a DDE-type transposase (Fig. S3), is the most abundant repeat in the *P. bonacinae* genome, followed by two similar satellite DNA candidates and Ty3-*gypsy* retrotransposons (Fig. 4J; the alignment of the satellite DNA candidates is shown in fig. S4). Ty3-*gyspy* retrotransposons, particularly a T2 element, predominate in the *P. americana* genome (Fig. 4H), and also contribute to the remaining *Prosopanche* species.

**Figure 3.**
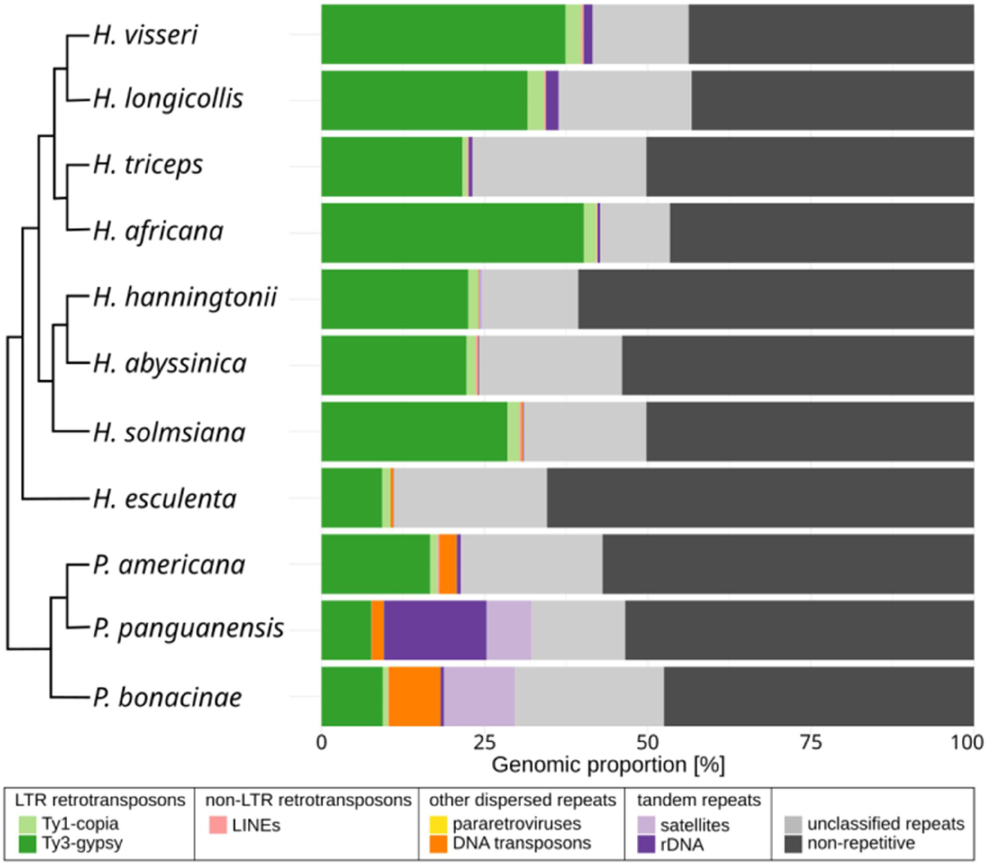
Repeat profiles of the analyzed Hydnoraceae genomes. The genomic abundance of each repetitive elements from the individual read clustering analysis using the RepeatExplorer2 pipeline were visualized as stacked bar charts (Tables S5 and S6). One bar indicates the entirety of one genome, where the stacks indicate the respective repeat composition. The bars for the respective species are arranged along the tips of the cladogram adapted from (Mkala *et al*., 2023).

**Figure 4.**
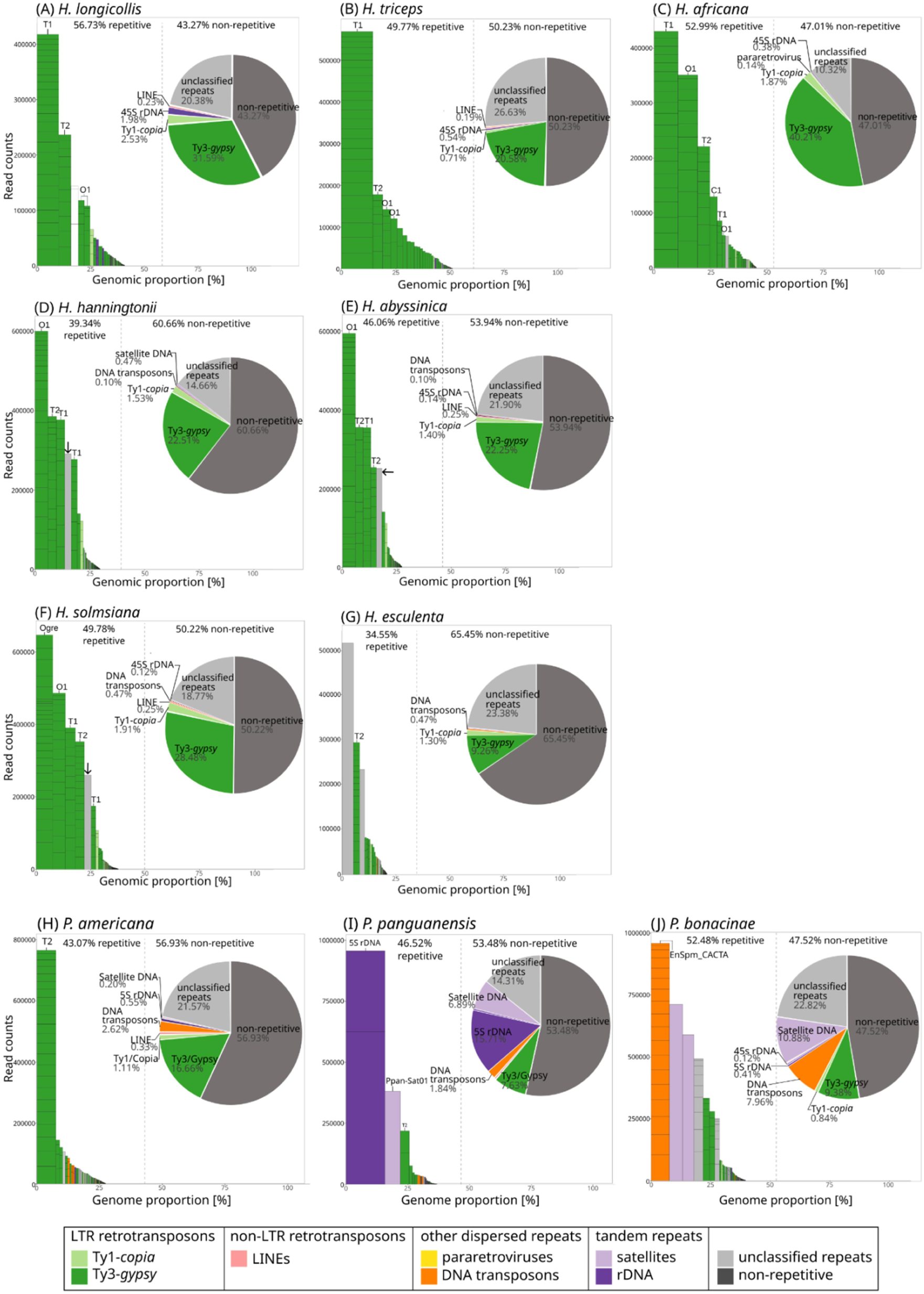
Repeat profiles of seven *Hydnora* and three *Prosopanche* genomes. The figures are arranged as (A-G): Repeat profiles of *Hydnora* genomes excluding aforementioned *H. visseri* genome; (H-J): Repeat profiles of *Prosopanche* genomes. Bars indicate the superclusters, with each stack representing a read cluster. The height and width of the bars correspond to the read counts within the supercluster and the genomic proportion, respectively. Superclusters were annotated using the reference repeat database of 17 reconstructed elements from the *H. visseri* genome. For *H. longicollis* (A), the third bar in the bar chart (colored in white) refers to a contamination of the input read dataset by short nucleotide sequences containing G- and A-stretches.

### Relative repeat abundances mirror the Hydnoraceae phylogeny

To provide a comparative overview of the repeat profiles of Hydnoraceae, subsets of reads from all studied Hydnoraceae species were pooled for a joint read clustering analysis. The read clusters were then annotated using the repeats database from the *H. visseri* genome, which allowed us assessing the presence of shared repeats across the analyzed Hydnoraceae genomes. Overall, the repeat profiles reflect the phylogenetic relationships: Phylogenetically close species share the same repeats (identified as shared protein domains) with similar genomic abundances (Figs 5 and 6). Taxa-specific repeats were observed as well, for instance, T4, O4, and O5 are exclusively observed in the genomes of *H. visseri*, *H. longicollis*, *H. triceps*, and *H. africana* (Fig. 6 and Table S7). *Hydnora esculenta*, being sister to the remaining *Hydnora* species, exhibits unique sequence variations within otherwise shared retrotransposons (e.g. T1 and T2; Fig. 5). Such sequence variations, reflected in the presence or absence of certain read clusters (indicated by black arrows in Fig. 5), are observed across all reconstructed repeats. *Prosopanche* repeat profiles vary to a higher degree compared to those of the sister genus *Hydnora*, particularly, but not only when examining the distribution of unclassified repeats (shown in gray in Fig. 5). The distribution of DNA transposons, 5S rDNA, and species-specific satellite DNAs characterizes the *Prosopanche* genomes, while *Hydnora* genomes show no contributions to the read clusters of these repeats (Fig. 6 and Table S7). The 5S rDNA clusters are composed almost entirely of reads from the *P. panguanensis* genome (Table S7), in contrast to the 35S ribosomal DNA, which are relatively evenly distributed and more abundant in some *Hydnora* species (Fig. 6 and Table S7). Finally, *Aristolochia fimbriata*, the photoautotrophic species closely related to Hydnoraceae, displays a completely distinct repeat profile (i.e. with Athila and Ogre-type retrotransposons (Ty3-*gypsy*) as most abundant transposable elements, with unique satellite DNAs, and many unclassified repeats; Fig. 5 and Table S7). These diverging repeat profiles of closely related taxa raise questions about their evolutionary path.

**Figure 5.**
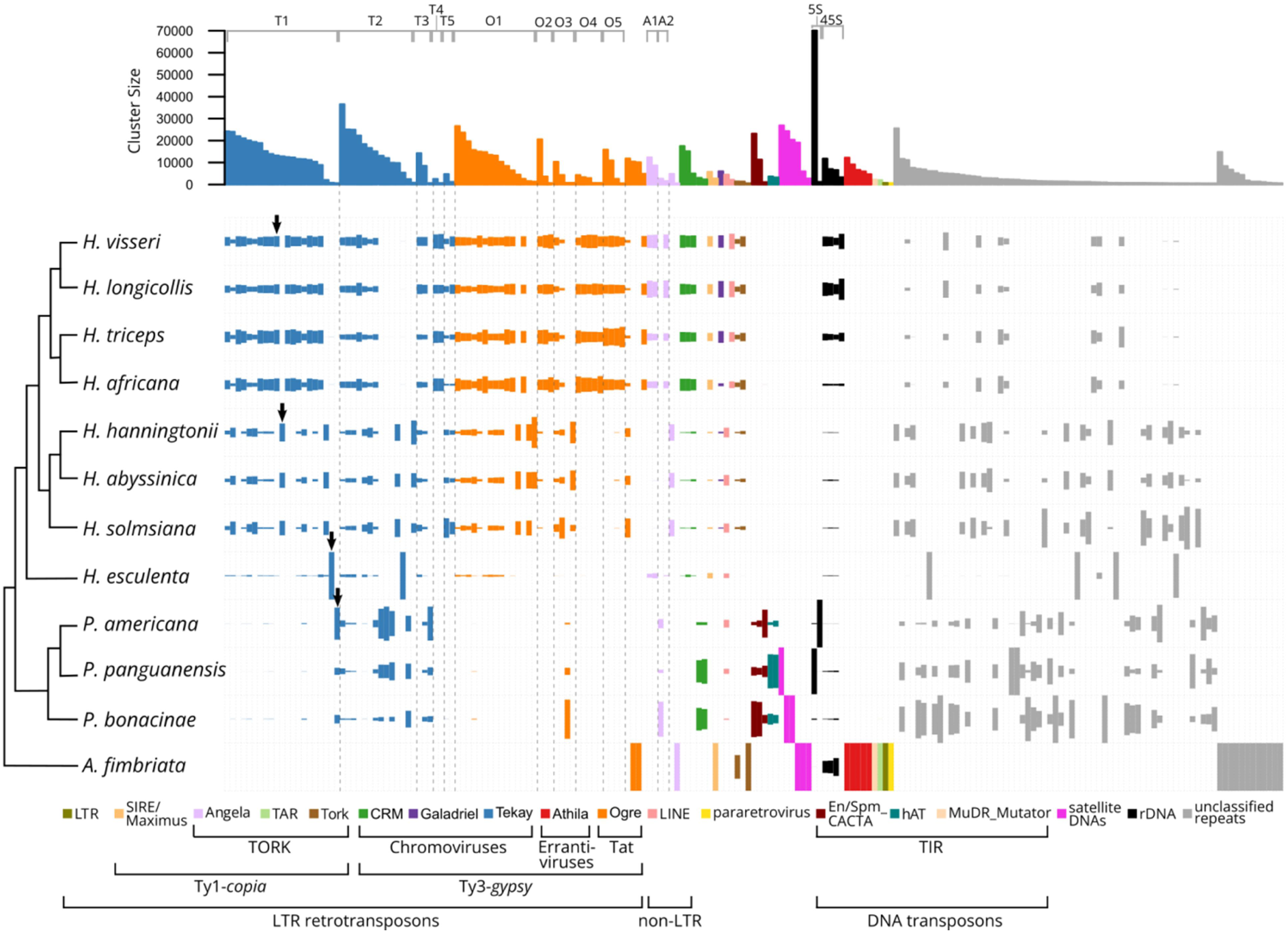
Comparative genomic repeat composition among eleven Hydnoraceae species and *A. fimbriata*. The size of each rectangle is proportional to the genomic abundance of the repeat in the respective species. Dashed vertical lines group read clusters that belong to the same repeat, while black arrows indicate taxa-specific sequence variants of *Hydnora*Tekay1. In the top bar chart, the height of each bar represents the number of reads in each cluster comprising at least 516 reads (≥ 0.01% of the analyzed reads). Clusters corresponding to the same repetitive element are sorted by read count, from highest to lowest, so the bar chart depicts the ranked abundance of each repetitive element. The phylogenetic tree is adapted from (Mkala *et al*., 2023). Please see Fig. S5 for a modified version of this figure, which includes the cluster identifiers.

**Figure 6.**
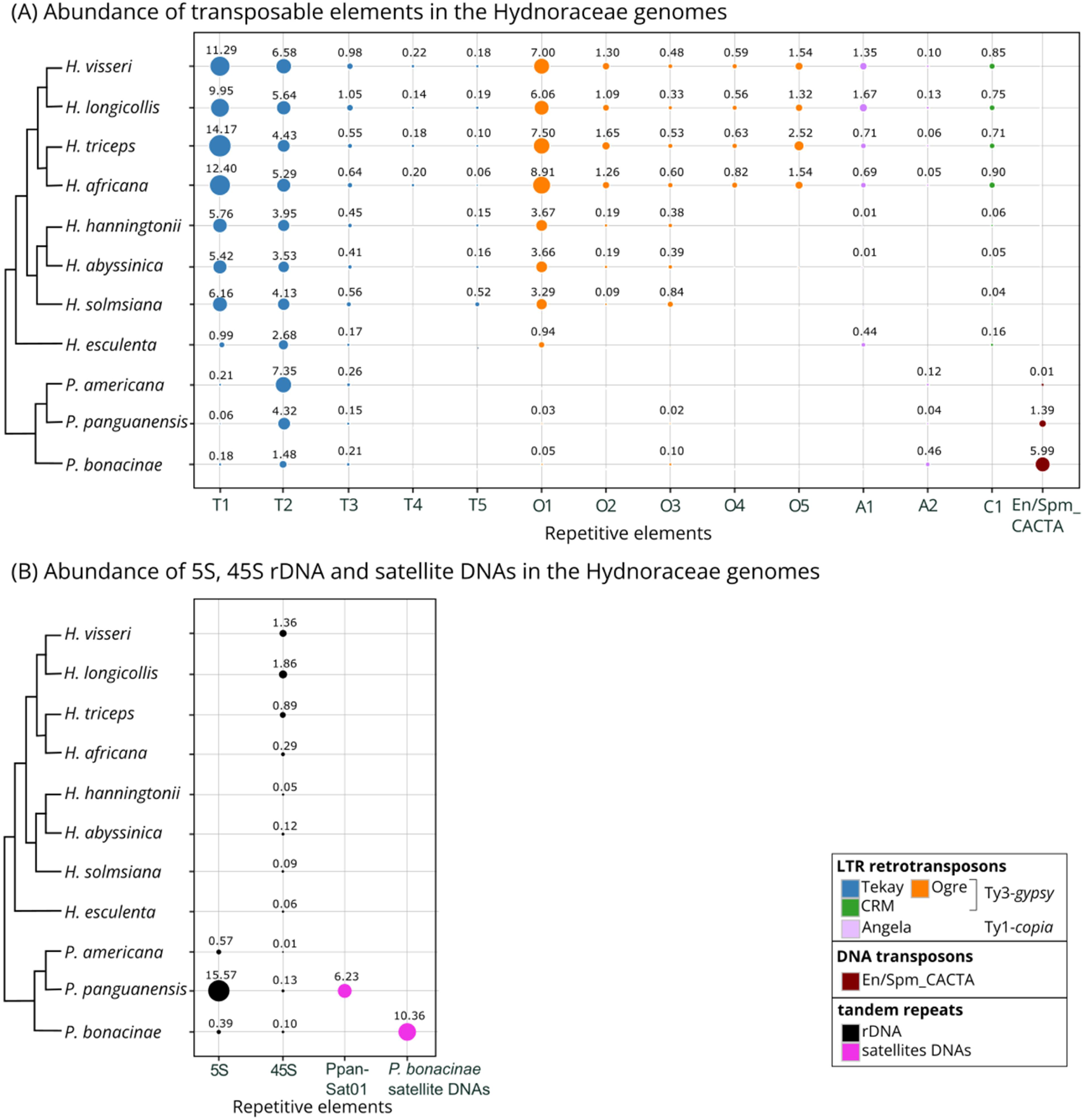
(A) The abundance of transposable elements in Hydnoraceae genomes. (B) 5S, 35S rDNAs, and satellite DNAs abundance in Hydnoraceae genomes. The size of each bubble indicates the relative abundance of the repeats among the analyzed species. This is calculated as the read counts of a repeat, specific to a species, divided by the total number of reads clustered for that repeat (see Table S7 for the proportions). The numbers above the bubble indicate the abundance of repeat within the species genome, calculated as the same read counts divided by total number of analyzed reads for that species, represented as percentage (%). The cladogram is adapted from (Mkala *et al*., 2023).

### Hydnoraceae 5S rDNA sequences vary greatly from those of other angiosperms

Among all analyzed *Hydnora* genomes, we detected the 5S rDNA only in the *H. triceps* genome, and in very low quantities (Table S3). This observation could result from the 5S rDNA copies being generally low in abundance in the *Hydnora* genomes, potentially falling below the detection threshold of the pipeline, or due to a high sequence divergence in 5S rDNAs that impede detection. In order to understand the low amount of 5S rDNA detected in *Hydnora* genomes, we reconstructed 5S rDNAs from all analyzed Hydnoraceae species using the 5S rDNA from the *H. triceps* genome. In comparison to an autotrophic relative, *Aritolochia fimbriata*, the total reads of *Hydnora* species mapped to the *H. triceps* 5S rDNA were considerably low (0.00005% - 0.003% versus 0.073%; Table S3). The reconstructed 5S rDNAs were then compared to those of other angiosperms using a genetic distance matrix, including the Piperales order (which contain Hydnoraceae) and the major host orders: Malpighiales and Fabales (Fig. 7). Genetic distances were calculated based on the 5S rDNAs sequence alignment across Hydnoraceae and 56 other angiosperms (Fig. 7, for alignment see fig. S6). With genetic distances greater than 0.15, indicating that over 15% of nucleotides were substituted, Hydnoraceae 5S sequences are largely divergent from those of other angiosperms (Fig. 7, for the values displayed on the heatmap, see Fig. S7). Specifically, genetic distances between Hydnoraceae 5S and closely related families within the Piperales, including Lactoridaceae, Aristolochiaceae, Saururaceae, and Piperaceae, exceed 0.16 (Figs 7 and S7). Furthermore, 5S rDNA from the major host lineages Malpighiales and Fabales show no particular similarity to Hydnoraceae 5S rDNAs, with genetic distances exceeding 0.19. Within Hydnoraceae, the genetic distance between *Hydnora* and *Prosopanche* 5S rDNAs ranges from 0.21 to 0.27, with 0.27 representing the highest genetic distance observed across all analyzed species. These variations occur throughout the DNA sequence, but are less pronounced within the internal control region (ICR) for transcription, specifically the A-box, Intermediate Element (IE), and C-box (Fig. S6). Overall, Hydnoraceae 5S are highly divergent from those of other angiosperms; however, since both *H. triceps* as well as *Prospanche* 5S rDNAs were successfully detected by the RE2 pipeline, sequence divergence does not account for the limited detection of the *Hydnora* 5S rDNA, but rather the low abundance of the 5S rDNA in *Hydnora* genomes.

**Figure 7.**
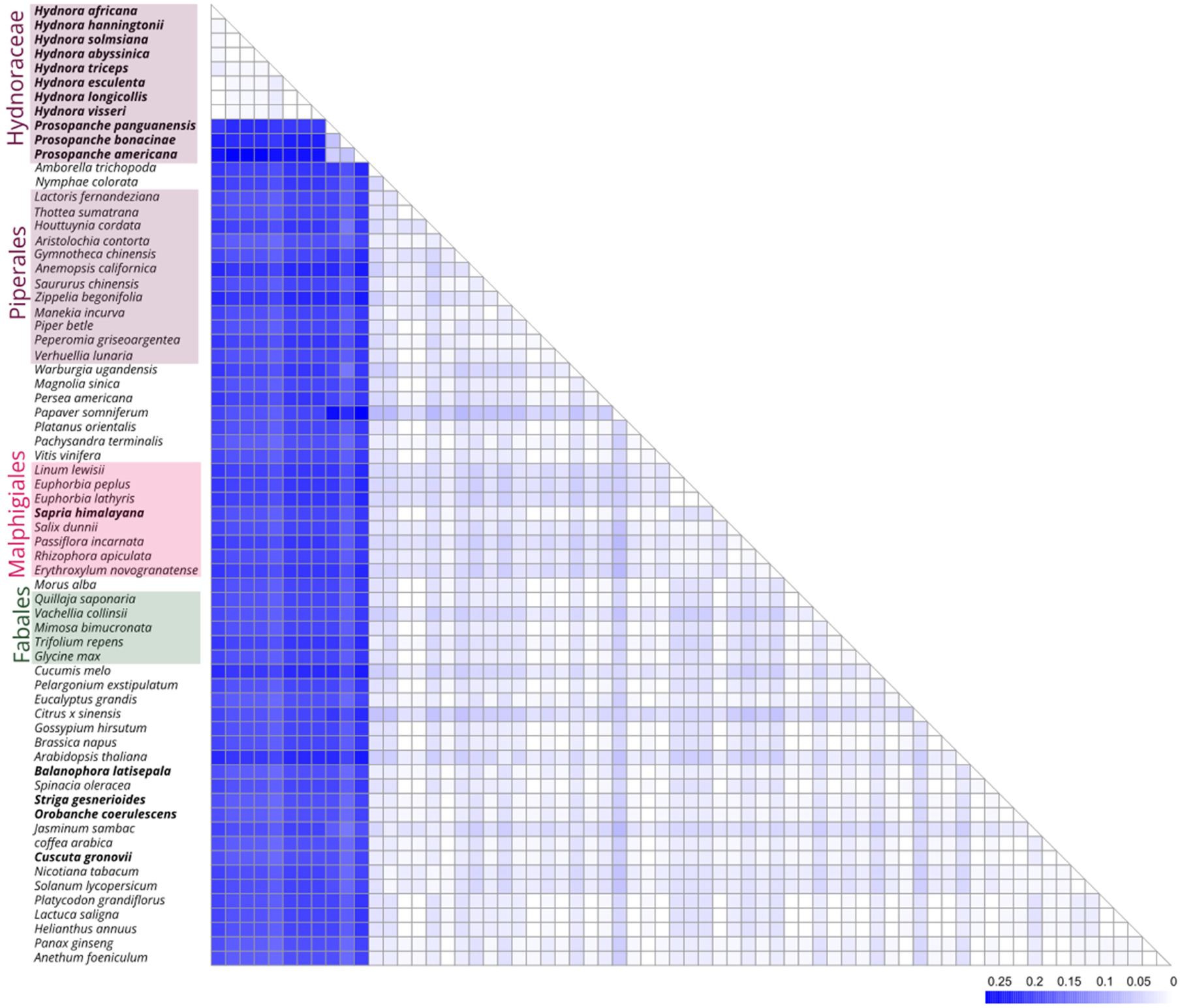
Genetic distances between Hydnoraceae 5S ribosomal DNAs and 56 other angiosperms, calculated using the Kimura-2-parameter model (Kimura, 1980). The resulting distance matrix ranges from 0 to 0.27, with higher values indicating greater sequence variation. Representatives of the order Piperales are highlighted in purple. Fabales and Malpighiales, the host orders, are highlighted in green and pink, respectively. Parasitic plants are highlighted in bold. Hydnoraceae, belonging to Piperales, are visualized on the top of the angiosperms to highlight the large genetic distances compared to other angiosperms.

### The 5S rDNA arrays in the *H. visseri* genome show a high degree of diversification

To capture the 5S rDNA organization within the *H. visseri* genome, a complete 5S rDNA monomer (121 bp gene and 977 bp intergenic spacer) was reconstructed and aligned to *H. visseri* long reads (Fig. 8). Among the 4,540,941 long reads analyzed (spanning approx. 500 bp up to 72,000 bp), 319 (0.007%) exhibit one or more aligned 5S rDNA monomer(s). These alignments vary greatly, which indicate diverging genomic organization of 5S rDNAs. Regular 5S rDNA arrays, consisting of at least five monomers arranged without major gaps in the reads longer than 6,000 bp, were identified 56 times (Fig. 8; A and A’). Some regular arrays exhibit sequence divergence or monomer rearrangements, such as insertions or overlapping monomers (Fig. 8A). Moreover, length-variants of the intergenic spacers were observed, recognizable by differing distances between the individual 5S rDNA monomers (Fig. 8A’, highlighted by arrows). The elongated intergenic spacers are due to internal sequence repetitions, as displayed in the dotplot (Fig. 8A’; Long ITS). Additionally, 24 potentially regular arrays were identified, however, due to read length limitations their integrity could not be confirmed. A total of 239 instances were identified as pseudogenic, characterized by large gaps between monomers due to sequences falling below the detection threshold (E-value = 0.0001; Fig. 8B). These pseudogenized arrays also showed high sequence variation, as indicated by faint lines within the dotplots, suggesting low 5S rDNA conservation in the *H. visseri* genome.

**Figure 8.**
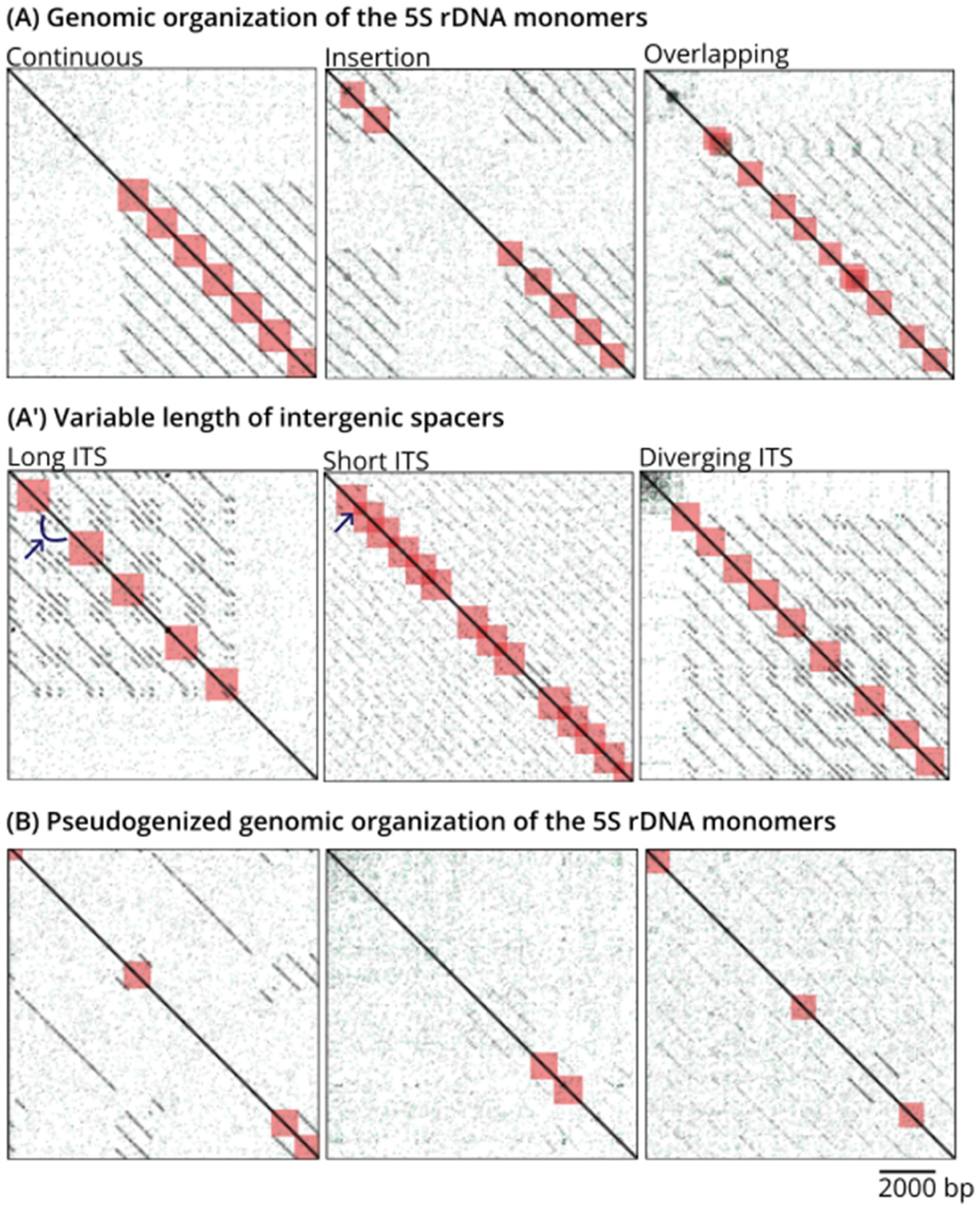
Genomic organization of *H. visseri* 5S rDNA monomers. Self-dotplots of PacBio long reads containing 5S rDNA arrays and monomers identified by BLAST (E-value = 0.0001) are displayed. Black lines parallel to the main diagonal represent sequence repetitions, while reversed sequence repetitions are displayed in green. Red boxes indicate a BLAST hit corresponding to the 5S rDNA gene (121 bp), including 500 bp up- and downstream of the gene (annotation zoom = 500). Thus, the tandem repetitions of the monomers include the intergenic spacers (977 bp) in between the genes. (A and A’) indicate regular arrangements of monomers, including length variants of intergenic spacers. The elongated and shortened ITS are highlighted by arrows. (B) displays potentially pseudogenized 5S rDNA arrangements, exhibiting large gaps between monomers on the faint, non-continuous lines in the dotplot.

## Discussion

To understand the genomic underpinning of parasitic plants, we focused on the repetitive DNAs recognizing them as one of the primary actors in genomic adaption to holoparasitic lifestyle. For the first time, we analyzed the repetitive genome content of eleven Hydnoraceae, ‘the strangest plants in the world’ (Musselman & Visser, 1989). Diverging repeatomes of Hydnoraceae are discussed under the hypothesis of rapid evolution, emphasizing the contrast between the two genera, *Hydnora* and *Prosopanche*. In *Hydnora*, we highlight the potential evolutionary roles of abundant LTR retrotransposons, whereas in *Prosopanche*, we observe a rather rare occurrence of very abundant 5S rDNA or DNA transposons, implying rapid evolution of repetitive fraction of the genome. Special focus lies on *Hydnora* 5S rDNAs, as eight *Hydnora* species almost completely lack 5S rDNA, which is a critical component for ribosomal biogenesis as well as chromosomal organization. Following the discussion on the individual repeatomes, the overall abundance of shared repeats is discussed in relation with the phylogeny, geographical distribution, and known host plant shifts, which underscore the role of repeats in shaping the genome and potential speciation.

### The diversified Hydnoraceae repeat landscape suggests rapid genome evolution

As in many seed plants, LTR retrotransposons are the main contributors to the *Hydnora* repeatomes (Wessler *et al*., 1995; Wells & Feschotte, 2020). Ty3-*gypsy* is the most abundant LTR retrotransposon superfamily across all *Hydnora* and *Prosopanche americana* genomes, while they rank second in the other two *Prosopanche* species analyzed. On average, Ty3-*gypsy* retrotransposons are 15.7 times more prevalent in the *Hydnora* genomes than those belonging to Ty1-*copia*, which is remarkable even taking into account that angiosperms generally contain more Ty3-*gypsy* (17–30%) compared to Ty1-*copia* retrotransposons (6–10%; Giraud *et al*., 2021). This large Ty3-*gypsy* fraction is dominated by a few high copy repeats, in particular two Tekay and one Ogre, suggesting their rapid accumulation in the recent past (Feschotte *et al*., 2002). The prevalence of Ty3-*gypsy* elements are worth investigating further in relation to their impact on genome size and chromosome configuration, e.g. chromosomal behavior in cell division and heterochromatin silencing, where Ty3-*gypsy* elements play a major role (Bourque *et al*., 2018; Orozco-Arias *et al*., 2019). Additionally, abundant transposable elements often correlate positively with genome size (Leeton & Smyth, 1993; Lee & Kim, 2014; Schmidt *et al*., 2024), also observed in parasitic plants (Neumann *et al*., 2021).

The reference repeat database generated based on the *H. visseri* genome effectively identifies repeats within the subgenus *Hydnora* (represented by *H. visseri*, *H. longicollis*, *H. triceps*, and *H. africana*). However, challenges remain when annotating closely related species. In particular, the *H. hanningtonii, H. abyssinica*, and *H. solmsiana* genomes contain highly abundant, nearly identical repeats that remain unclassified. Likewise, two major clusters in the *H. esculenta* genome are yet unclassified. These repeat identification gaps suggest that neither the *H. visseri* reference repeat database nor the pipeline’s internal database adequately captures the repeat diversity in Hydnoraceae, especially outside subgenus *Hydnora*. Several factors contribute to the gaps: Firstly, apart from those presented here, there are no repetitive DNA sequences known for Hydnoraceae. Second, parasitic plant genomes are subject to distinct evolutionary pressure compared to photoautotrophic plants due to their parasitic lifestyle, represented as large scale gene losses or accelerated evolutionary rate (Bromham *et al*., 2013; Sun *et al*., 2018; Lyko & Wicke, 2021; Xu *et al*., 2022; Ashapkin *et al*., 2023; Chen *et al*., 2023). Furthermore, repetitive DNAs tend to evolve faster than the coding regions of genes due to reduced selection pressure (Mehrotra & Goyal, 2014). In summary, Hydnoraceae repeatomes may have undergone rapid evolution under reduced selective pressure.

The assumed rapid evolution in Hydnoraceae repeatomes aligns with the repeat variability observed in the *Prosopanche* genomes, which revealed very different repeat profiles compared to *Hydnora*. In particular, the high 5S rDNA abundance in *P. panguanensis* as well as the high En/Spm_CACTA (DNA transposon) abundance in *P. bonacinae* deserve further attention. 5S rDNA being the most abundant repetitive element in a genome is to our knowledge unprecedented, and could impede chromosomal stability via frequent recombination (Raskina *et al*., 2008; Ding *et al*., 2022). Similarly, DNA transposons are usually less abundant in plant genomes compared to LTR retrotransposons (Feschotte & Pritham, 2007), due to several factors like silencing by the host genome, cut-and-paste transposition (in comparison to the copy- and-paste transposition of retrotransposons), and hindrance from retrotransposons which might target the catalytic region of DNA transposons (Zhao *et al*., 2016; Liu *et al*., 2021). However, active En/Spm_CACTA elements could transpose into the proximity of the coding regions of genes and can play a major role in genome divergence as observed in rice, cabbage, and maize (Jiang *et al*., 2003; Alix *et al*., 2008; Yang *et al*., 2013). In addition, rapid evolution through actively amplifying DNA transposons has been observed occasionally across the tree of life, e.g. within genomes of taxa with rapid speciation records such as bats (Mitra *et al*., 2013; Paulat *et al*., 2022). Thus, the abundant En/Spm_CACTA in the *P. bonacinae* genome is worth investigating further in terms of a potential repeat-driven evolution.

Summarizing, a fast evolution of Hydnoraceae genome is supported by (1) the predominance of certain repeats in the genome, for instance, DNA transposons in *P. bonacinae*; and (2) the overall repeat diversity across the Hydnoraceae genomes. However, to confirm the potentially accelerated evolutionary rate in the Hydnoraceae nuclear genome, further validation, e.g. by comparing the divergence of orthologous single-copy genes is required. Moreover, it will be necessary to also compare against the genomes of closely related photoautotrophic species, as well as against genomes of different parasitic lineages.

### Atypical 5S rDNA abundance, sequence variation, and genomic organization challenge our current understanding

The scarcity of 5S rDNA in *Hydnora* genomes sharply contrasts with the large numbers of 5S rDNA copies typically observed in autotrophic plants, which can range from a few hundred to more than 100,000 copies for a haploid genome (Danna *et al*., 1996; Fulneček *et al*., 2002; Wang *et al*., 2019). Since 5S rDNA plays a crucial role in maintaining chromosomal structure and supporting ribosome assembly, such a low abundance is likely to pose challenges for both chromosomal stabilization and rRNA transcription (Moss, 2004; Raskina *et al*., 2008; Kobayashi, 2011; Volkov *et al*., 2022). Additionally, *Hydnora* 5S rDNA exhibits high sequence variation compared to that of other angiosperms, including its autotrophic relatives and host plant families. Such sequence variation potentially influences the functionality of the 5S rDNA, as maintaining the secondary and tertiary structure is crucial for the RNA-protein interaction during ribosome assembly (Ciganda & Williams, 2011). Moreover, the genomic organization of the 5S rDNA in *H. visseri* is highly irregular, with sporadic and discontinuous monomer repetitions that putatively represent remnants of once larger, more cohesive arrays. The observed fragmentation and re-organization alongside the nucleotide sequence variation, and the overall scarcity of 5S rDNA in the *Hydnora* genomes, suggest a lack of conservation, possibly reflecting a weakened selective pressure on the 5S gene, similar to the accelerated molecular evolutionary rate seen in the Hydnoraceae 18S rDNA (Bromham *et al*., 2013). Although holoparasitic plants experience a reduced selective pressure and loss of many gene families (Vogel *et al*., 2018; Sun *et al*., 2018; Chen *et al*., 2023), a reduced selective pressure on rRNA-encoding genes is notable due to their importance for basic cellular machineries, such as ribosomes.

However, further investigation is required to determine whether the low genomic abundance and sequence heterogeneity necessarily lead to a reduced or non-functionality, as the promoter regions may remain relatively intact, and deviating secondary structures may be tolerated as recent studies suggest that ribosomal DNA sequence heterogeneity may be more common in eukaryotes than previously thought (Wang *et al*., 2023). The observed 5S rDNA heterogeneity in Hydnoraceae, may be the result of ‘incomplete’ concerted evolution that could be caused by a variety of factors, including genome dynamics (e.g. whole genome duplication, chromosome aberrations, and activity of transposable elements), life history traits (e.g. long lifespan, asexual reproduction), and severe environment (Wang *et al*., 2023; Garcia *et al*., 2024). This suggests that the evolution of *Hydnora* 5S rDNA is possibly linked to the large-scale genome alterations, which may be related to the parasitic lifestyle (Sun *et al*., 2018; Chen *et al*., 2023).

Several key questions remain unanswered: As the genome size of these plants are not yet known to science, what would be the absolute 5S rDNA amount in *Hydnora* species? If *Hydnora* genomes were as big as assumed (>10 GB, dePamphilis, personal communication), would the small 5S rDNA copy numbers be sufficient to be functional in the cell? Are *Hydnora* ribosomes actually composed of 5S rRNA with an alternate sequence, possibly benefiting from the rRNA/ribosome of their host (Yang *et al*., 2019)? Understanding the genome size and identifying the 5S and 35S rDNA chromosomal loci would provide an opportunity for putting the findings of this study into a broader evolutionary context.

### Hydnoraceae repeat profiles reflect their phylogeny, geographic distribution, and host shifts

The repeat landscape across the analyzed Hydnoraceae species is largely congruent with a recent plastid-based Hydnoraceae phylogeny (Mkala *et al*., 2023), mirrored by a similar repeat composition and the individual repeat abundances in closely related species as well as the presence of (sub-)genus-specific repeats. The species of the subgenus *Hydnora* (former *Euhydnora*, represented here by *H. visseri*, *H. longicollis*, *H. triceps*, and *H. africana*; (Decaisse, 1873; Hatt *et al*., 2024b)) share the same Tekay element as their most abundant repeat (T1) and harbor several clade-specific elements (T4, O4, and O5). These species are exclusively found in southern Africa (mostly South Africa, Namibia, and southern Angola) and parasitize *Euphorbia* spp. (Decaisse, 1873; Musselman & Visser, 1989; Hatt *et al*., 2024b). In contrast, the species of the subgenus *Dorhyna* (represented here by *H. hanningtonii*, *H. abyssinica,* and *H. solmsiana*) are distributed in eastern and northern Africa, including the Arabian peninsula, and parasitize Fabaceae spp. (Decaisse, 1873; Musselman & Visser, 1989; Hatt *et al*., 2024b). These differences in lifestyle are accompanied by differences in the repeat profile, such as the predominance of Ogre instead of Tekay elements and the presence of further clade-specific repeats. *Hydnora esculenta,* endemic to Madagascar and the sole member of the subgenus *Neohydnora* (Harms, 1935; Bolin & Musselman, 2013), exhibits a unique repeat profile, i.e. lacking most of the Tekay and Ogre retrotransposons found in other *Hydnora* species, but also harbor neither identifiable DNA transposons nor identifiable satellite DNAs. This contrasts with the other analyzed *Hydnora* and *Prosopanche* genomes, resulting in an overall low repeat content in the *H. esculenta* genome. While the other *Hydnora* species’ habitat is in arid and semi-arid climates, *H. esculenta* is also found in transitional rainforest areas (Bolin & Musselman, 2013; Hatt *et al*., 2024b). The distinct growing features indicate that the species has undergone genomic adaptation to distinct environmental stress, which may be reflected in the repeat profile, via genetic drift and/or reduced gene flow due to population isolation, but also stress-induced transposable element activation (Casacuberta & González, 2013). Emerging questions include the possible impact of horizontal repeat transfers from host to parasite and how these might shape *Hydnora* genomes (Baidouri *et al*., 2014).

In contrast to *Hydnora*, the geographic distribution of *Prosopanche* species does not closely align with their phylogeny. For instance, although *P. americana* and *P. panguanensis* are phylogenetically closer, *P. americana* and *P. bonacinae* are sympatrically distributed in the arid regions of Argentina, while *P. panguanensis* inhabits the tropical rainforest of Peru (Hatt *et al*., 2023). Given that only 40% of known *Prosopanche* species were analyzed in this study, compared to 80% of the known *Hydnora* species diversity, the limited sampling may restrict a more comprehensive analysis, particularly as new species are likely to be discovered (Hatt *et al*., 2023). In addition, considering that most Hydnoraceae species are found in semi-arid climates, the habitat of *P. panguanensis* in rainforest is again noticeable (Martel *et al*., 2018). The unique, massive 5S rDNA in the *P. panguanensis* genome might thus be related to the habitat, and may have contributed to the adaptation capacity of the species, since rDNA is known to represent recombinogenic ‘hot spots’, and could thus lead to speciation by chromosomal rearrangement, when combined with geographical isolation and establishment by genetic drift (Raskina *et al*., 2008; Rosato *et al*., 2015).

Together, out study marks the evolutionary role of repetitive DNAs in holoparasitic plant genome, contributing to fill the large gap in genomic research of holoparasitic plants. The diverging repeat profiles, i.e. the high LTR retrotransposon content in *Hydnora* in contrast to the abundant 5S rDNA, DNA transposons, and satellite DNAs in *Prosopanche*, alongside with the unclassified, abundant repeats in some *Hydnora* genomes, suggest a relatively unrestrained repeatome evolution from selective pressure. The variable 5S rDNA sequences in Hydnoraceae and their fluctuating abundance between the genomes further challenge the 5S rDNA functionality in Hydnoraceae genomes, underscoring a potentially rapid evolution. For a more comprehensive decoding of the evolutionary pattern, we propose a possible link between geographical distribution, host shifts, and repeat compositions of Hydnoraceae, hypothesizing that the accumulation patterns of repetitive elements contribute to genomic adaptation. Despite the so-far limited nuclear genomic research on parasitic plants, our findings align with studies on other parasitic plant taxa such as Balanophoraceae, Orobanchaceae, and *Cuscuta*, which also experience reduced selective pressure (Xu *et al*., 2022; Ashapkin *et al*., 2023; Chen *et al*., 2023). However, we also want to pinpoint the limitations of our study, such as the scarcity of materials due to absence of cultivation, the lack of material for genome size measurements, and hence a potential underestimation of repeats due to thresholds in the repeat clustering procedure. Nevertheless, these gaps underscore the significance of the present study, adding valuable insights into the evolution of parasitic plant nuclear genomes in general.

## Acknowledgements

We would like to thank Andrea Coccuci and Jay F. Bolin for providing us with additional photos for *Hydnora* and *Prosopanche* and for giving their consent for these photos to be published. Additionally, we would like to thank Andreas Dahl (Dresden Concept Genome Center) for his support in sequencing of Hydnoraceae. We thank Claude dePamphilis (Huck Institute of the Life Science, Penn State University) for the discussion regarding the genome size of Hydnoraceae. We thank Sònia Garcia (CSIC, Institut Botànic de Barcelona) for the many discussions regarding the 5S rDNA in the Hydnoraceae.

## Author contributions

St.Wa., T.H. conceived the study; data generation W.K., N.S., E.M.M., G.W.H., M.J., St.Wa., Sy.Wi.; analyses W.K., N.S.; writing of the first draft W.K., N.S; responsible for visualization of results W.K., N.S.; all authors reviewed and edited the draft and agreed to the published version of the manuscript.

## Competing interests

The authors declare that they have no competing interests.

## Data availability

The dataset generate and/or analyzed during the current study are available in the public data repositories under the identifier: PRJNA1188088 (BioProject, NCBI genebank) and https://doi.org/10.5281/zenodo.14194251 (Zenodo).

The sequence data are stored in the public data repository under the BioProject PRJNA1188088 (NCBI genebank) and https://doi.org/10.5281/zenodo.14194251 (Zenodo). This information is included in the paper under the section ‘Data availability’.

## Supporting Information

The supporting information for the current study contains

**Fig. S1** Workflow for the processing of the read data (A) and the reconstruction of a preliminary reference database of transposable elements within the *H. visseri* genome (B)

**Fig. S2** Alignment of the consensuses of the highly abundant unclassified read cluster from the *H. abyssinica*, *H. hanningtonii*, and *H. solmsiana* individual analysis results

**Fig. S3** The reconstructed consensus of *P. bonacinae* En/Spm_CACTA transposase aligned with transposase amino acid sequences from 16 further En/Spm_CACTA DNA transposon sequences

**Fig. S4** Alignment of the RE2-provided consensuses from two highly abundant read clusters representing *P. bonacinae*-specific satellite DNAs

**Fig. S5** Comparative genomic repeat composition among eleven Hydnoraceae spp. and *A. fimbriata*, including the Cluster IDs

**Fig. S6** Alignment of Hydnoraceae 5S rDNAs with 56 angiosperms 5S rDNAs

**Fig. S7** Genetic distances between Hydnoraceae 5S ribosomal DNAs and 64 other angiosperms, calculated using the Kimura-2-parameter model, including the genetic distances

**Table S1** Plant material, DNA extraction and genome sequencing

**Table S2** Reconstruction of repetitive elements within the *H. visseri* genome

**Table S3** Hydnoraceae short read mapping to 5S rDNA references

**Table S4** NCBI identifiers for the input genome sequencing data and the reconstructed 5S rDNAs for 5S rDNA sequence comparison between Hydnoraceae and angiosperms (Separated Excel file is provided)

**Table S5** Summary of genomic proportions of repetitive elements in *Hydnora* genomes

**Table S6** Summary of genomic proportions of repetitive elements in *Prosopanche* genomes

**Table S7** Relative genomic abundance of abundant repeats in Hydnoraceae genomes

## Notes

https://zenodo.org/records/14194251

### Competing Interest Statement

The authors have declared no competing interest.

