## Supplementary material for "Diverging repeatomes in holoparasitic Hydnoraceae uncover a playground of genome evolution": Supplimentary Figure 1-7 and Supplimentary Table 1-7

### *New Phytologist* Supporting Information

Article acceptance date: Click here to enter a date.

The following Supporting Information is available for this article:

**Fig. S1** Workflow for the processing of the read data (A) and the reconstruction of a preliminary reference database of transposable elements within the *H. visseri* genome (B)

**Fig. S2** Alignment of the consensuses of the highly abundant unclassified read cluster from the *H. abyssinica*, *H. hanningtonii*, and *H. solmsiana* individual analysis results

**Fig. S3** The reconstructed consensus of *P. bonacinae* En/Spm_CACTA transposase aligned with transposase amino acid sequences from 16 further En/Spm_CACTA DNA transposon sequences

**Fig. S4** Alignment of the RE2-provided consensuses from two highly abundant read clusters representing *P. bonacinae*-specific satellite DNAs

**Fig. S5** Comparative genomic repeat composition among eleven Hydnoraceae spp. and *A. fimbriata*, including the Cluster IDs

**Fig. S6** Alignment of Hydnoraceae 5S rDNAs with 56 angiosperms 5S rDNAs

**Fig. S7** Genetic distances between Hydnoraceae 5S ribosomal DNAs and 64 other angiosperms, calculated using the Kimura-2-parameter model, including the genetic distances

**Table S1** Plant material, DNA extraction and genome sequencing

**Table S2** Reconstruction of repetitive elements within the *H. visseri* genome

**Table S3** Hydnoraceae short read mapping to 5S rDNA references

**Table S4** NCBI identifiers for the input genome sequencing data and the reconstructed 5S rDNAs for 5S rDNA sequence comparison between Hydnoraceae and angiosperms (Separated Excel file is provided)

**Table S5** Summary of genomic proportions of repetitive elements in *Hydnora* genomes

**Table S6** Summary of genomic proportions of repetitive elements in *Prosopanche* genomes

**Table S7** Relative genomic abundance of abundant repeats in Hydnoraceae genomes

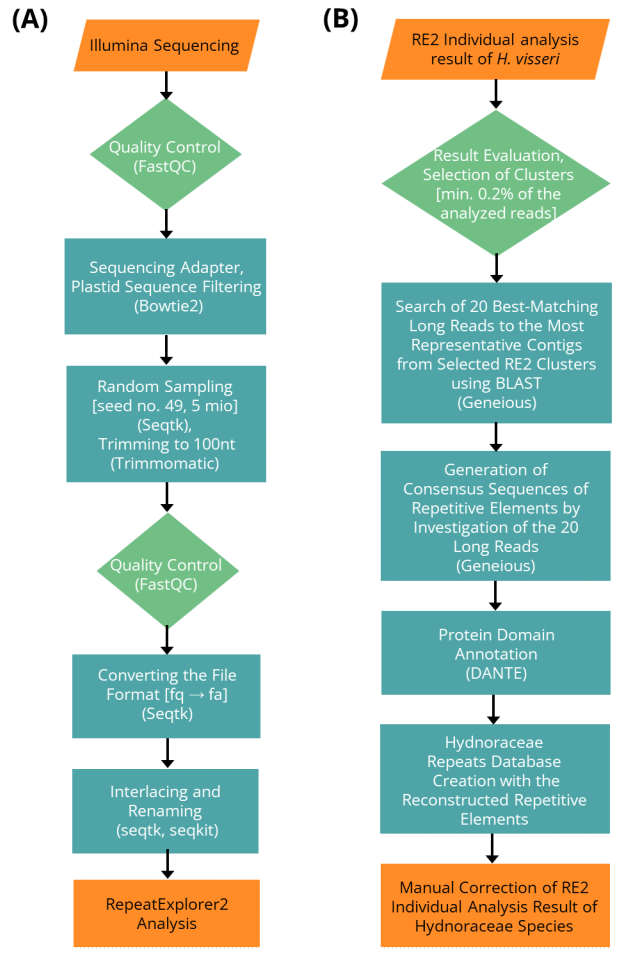

**Fig. S1** Workflow for the processing of the read data (A) and the reconstruction of a preliminary reference database of transposable elements within the *H. visseri* genome (B). Parallelograms indicate input data, whereas rectangles and hexagons indicate processes and preparation steps, respectively. Light green rectangles indicate processes in which interim results were generated. (A) In order to be usable in the RepeatExplorer2 (RE2) analysis, the Illumina reads must meet certain requirements, such as having a certain length and being provided in a certain file format. (B) The RE2 results (graph-based clusters) of the analysis using *H. visseri* read data were used for the generation of a custom database containing the most abundant repeats within the *H. visseri* genome.

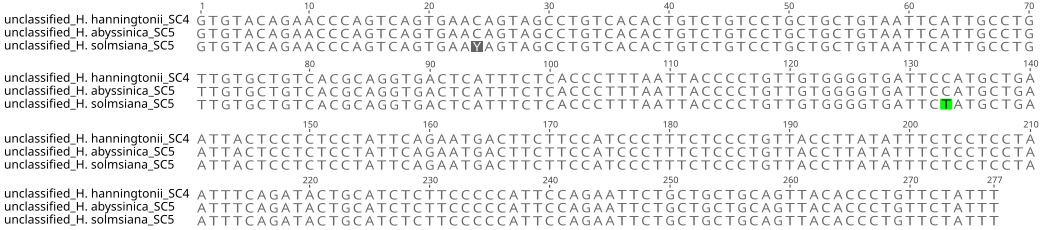

**Fig. S2** Alignment of the consensuses of the highly abundant unclassified read cluster from the *H. abyssinica*, *H. hanningtonii*, and *H. solmsiana* individual analysis results. The contigs from the specific RE2 cluster were aligned to the highest read depth contig from the cluster to refine the consensuses. The repeat consensuses have the same length of 277 bp, and one nucleotide substitution and one potential nucleotide substitution were observed from the consensuses alignment. The repeat consensuses could neither be identified using the RE2 database, nor by publicly available databases.

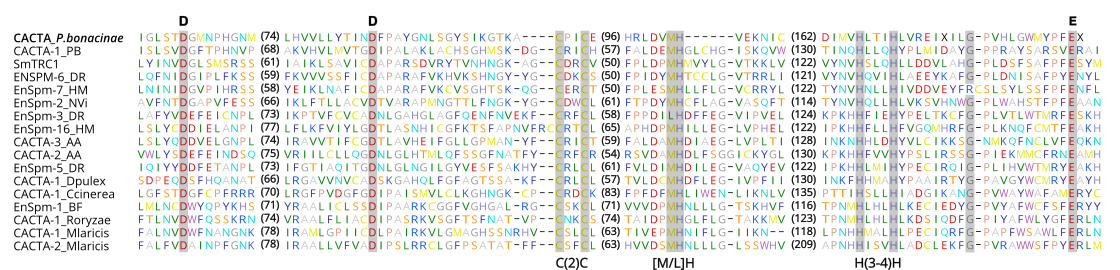

**Fig. S3** The reconstructed consensus of *P. bonacinae* En/Spm_CACTA transposase aligned with transposase amino acid sequences from 16 further En/Spm_CACTA DNA transposon sequences, which represent this transposon superfamily, from (Yuan & Wessler, 2011). The pivotal cataylic region represented by three amino acid (D, D, E) and the additional conserved motifs, C(2)C, [M/L]H, and H(3-4)H are shaded.

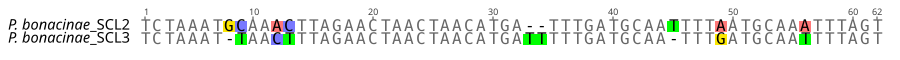

**Fig. S4** Alignment of the RE2-provided consensuses from two highly abundant read clusters representing *P. bonacinae*-specific satellite DNAs. In the *P. bonacinae* individual analysis, two highly abundant read clusters (supercluster 2 and 3) were identified as satellite DNAs. The alignment revealed the same length (60 bp) of both consensuses, potentially originating from a single ancestral sequence, diverged due to nucleotide substitutions as well as potential insertion/deletion mutations.

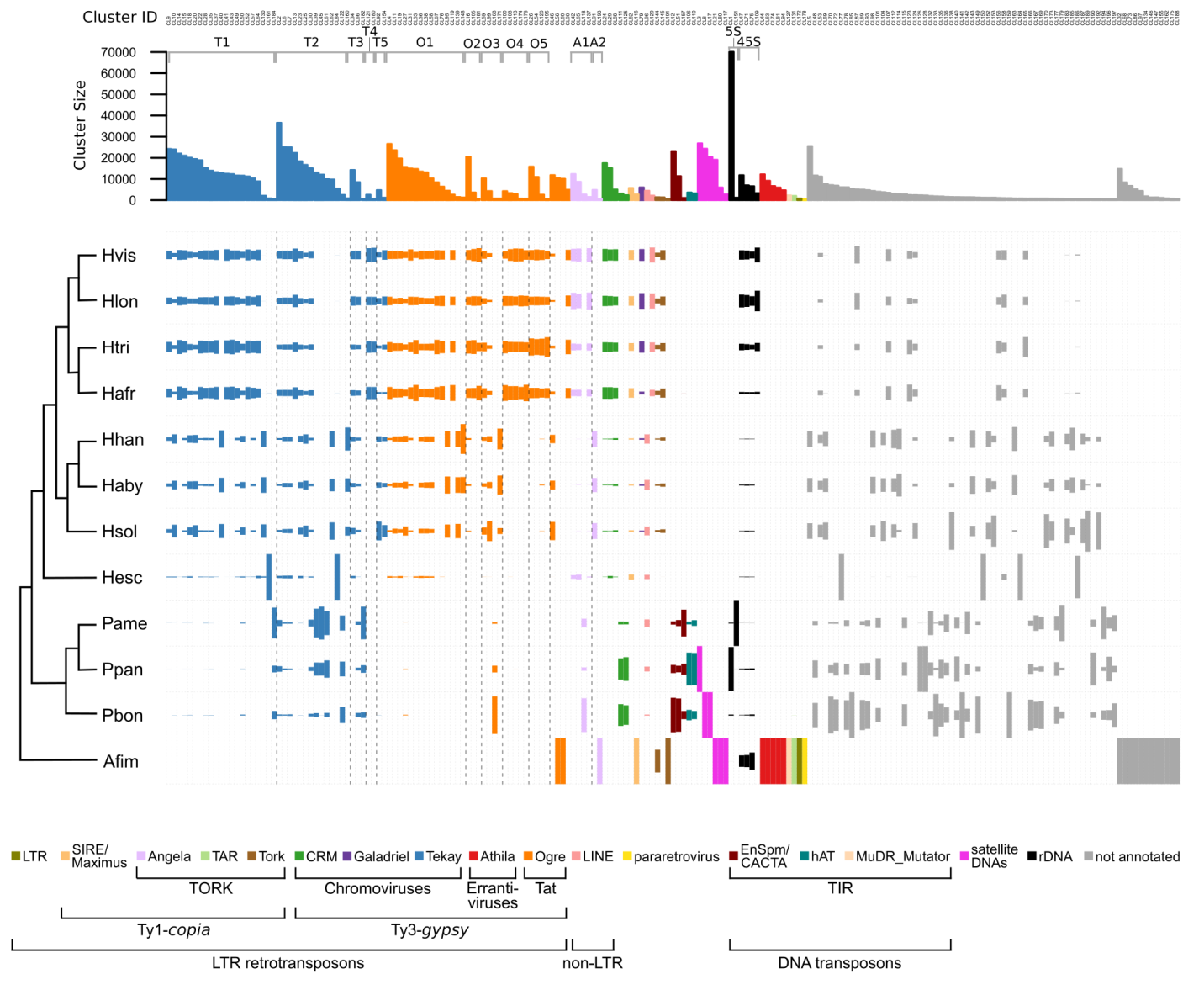

**Fig. S5** Comparative genomic repeat composition among eleven Hydnoraceae spp. and *A. fimbriata*, including the Cluster IDs (For legend, see Fig. 5)

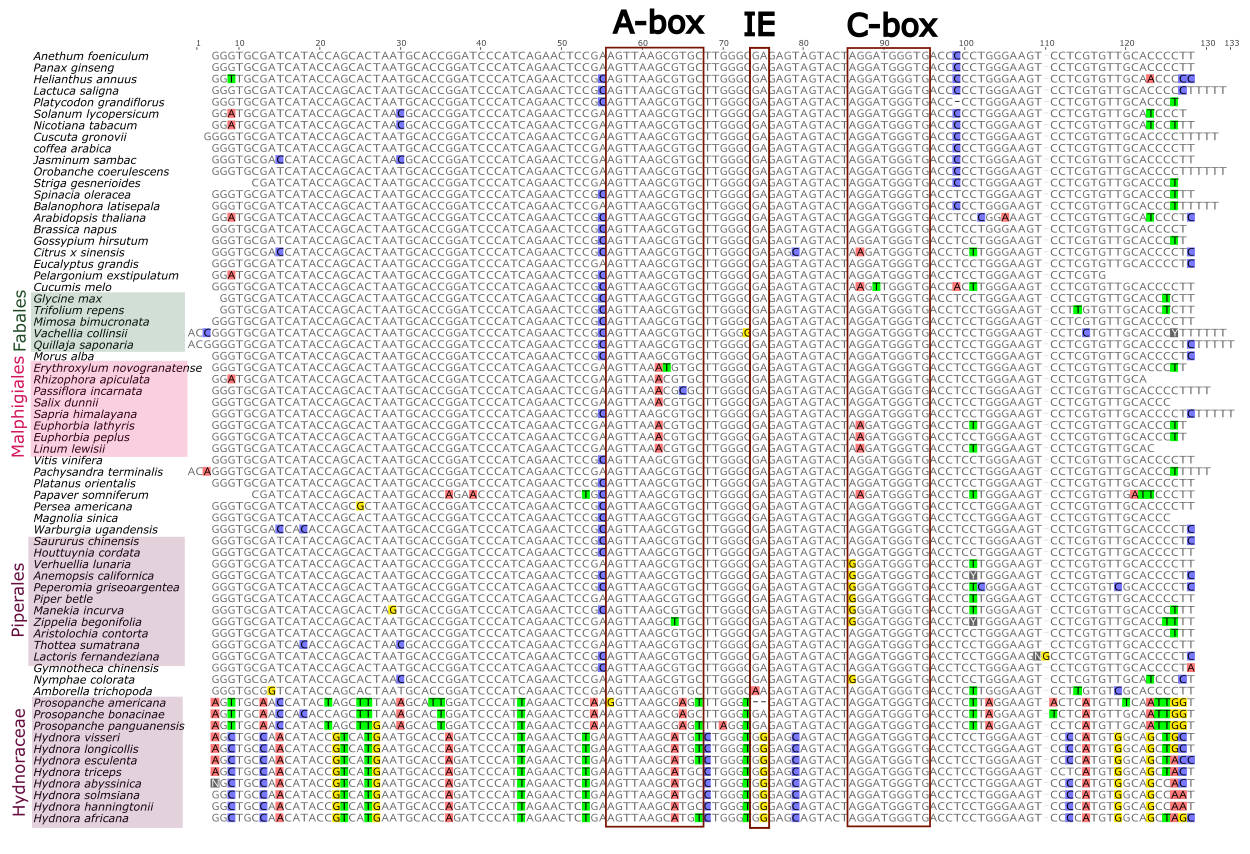

**Fig. S6** Alignment of 5S rDNAs from the Hydnoraceae and from 56 further angiosperms. Disagreements from the overall consensus sequence are highlighted (A = red, T = green, C = blue, G = yellow). The highly conserved internal control region (ICR) for transcription (A-box, Intermediate element=IE, C-box) is indicated in red boxes referring to (Maiwald *et al.*, 2024)

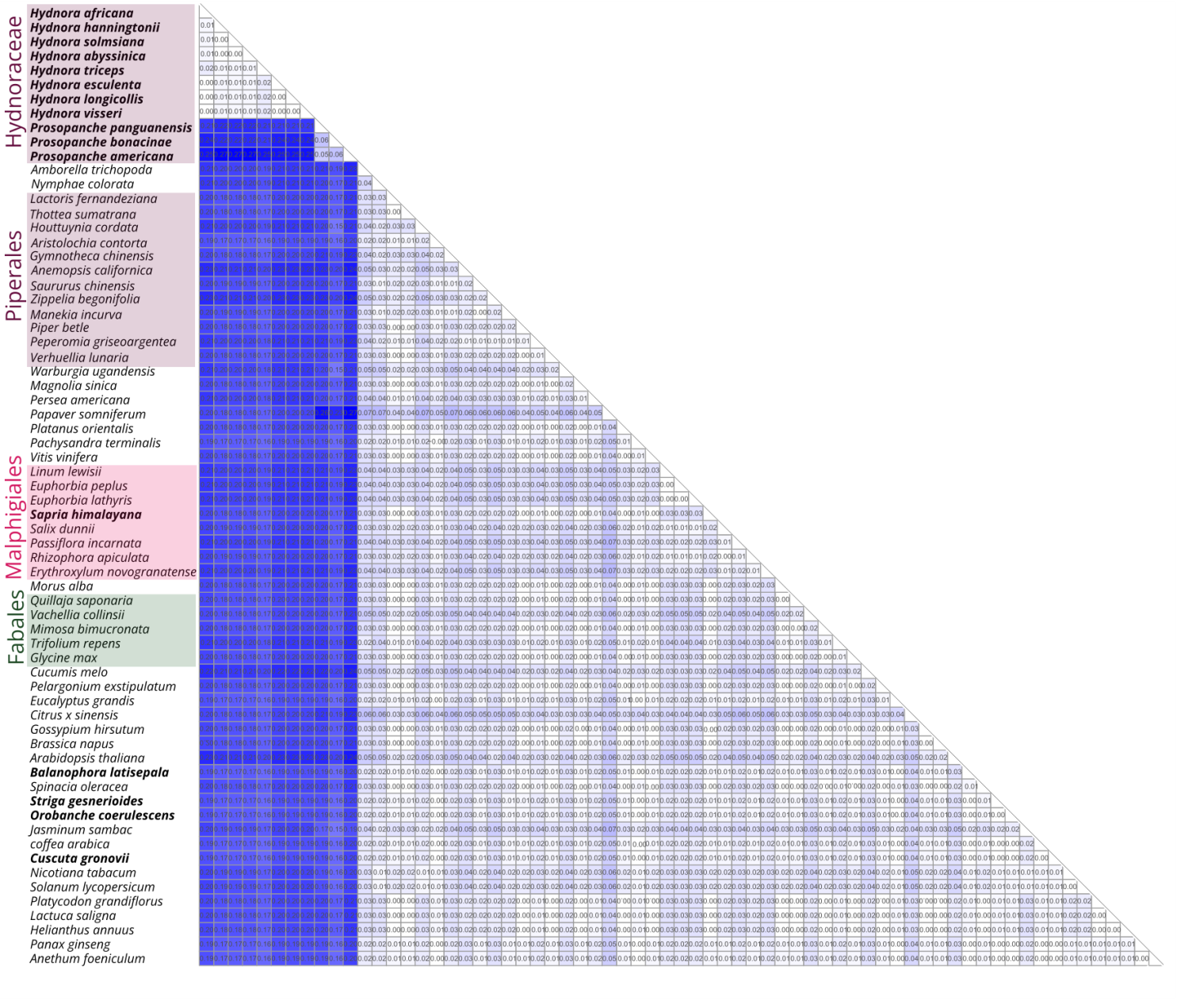

**Fig. S7** Genetic distances between the 5S ribosomal DNAs of the Hydnoraceae and 56 further angiosperms, calculated using the Kimura-2-parameter model (distances are shown) (for legend, see Fig. 7).

**Table S1** Plant material, DNA extraction and genome sequencing

| Taxon | Taxon Codes | DNA extraction method | Library preparation | Sequencing platform | Read length | Reference | NCBI identifiers |
| --- | --- | --- | --- | --- | --- | --- | --- |
| *Hydnora visseri* | Hvis | DNeasy Plant Mini Kit (Qiagen) | NEBNext Ultra DNA | Illumina HiSeq-2000 | 100 | (Naumann *et al.*, 2016) | SAMN44848561 |
|  |  | Modified (Doyle & Doyle, 1987) adding RNAse A (Thermo Scientific, Waltham, MA, USA) treatment (10 mg/ml) | BluePippin™ Size-Selection System | 43 PacBio RS2 SMRT cells v3 | 500 – 71,657 | This study |  |
| *Hydnora africana* | Hafr | Modified (Doyle & Doyle, 1987) adding RNAse A (Thermo Scientific, Waltham, MA, USA) treatment (10 mg/ml) | NEBNext Ultra DNA | Illumina HiSeq 2500 RapidMode | 150 | (Jost *et al.*, 2022) | SAMN44848557 |
| *Hydnora longicollis* | Hlon | DNeasy Plant Maxi Kit (Qiagen) | NEBNext Ultra DNA | Illumina HiSeq 2500 RapidMode | 150 | (Jost *et al.*, 2022) | SAMN44848559 |
| *Hydnora triceps* | Htri | Modified (Doyle & Doyle, 1987) adding RNAse A (Thermo Scientific, Waltham, MA, USA) treatment (10 mg/ml) | NEBNext Ultra DNA | Illumina HiSeq 2500 RapidMode | 150 | (Jost *et al.*, 2022) | SAMN44848560 |
| *Hydnora abyssinica* | Haby |  | Illumina TruSeq DNA | Illumina HiSeq-2000 | 150 | (Mkala *et al.*, 2023) | SAMN44821594 |
| *Hydnora hanningtonii* | Hhan |  | NEBNext Ultra DNA | Illumina HiSeq 2500 RapidMode | 150 | (Jost *et al.*, 2022) |  |
| *Hydnora solmsiana* | Hsol |  | NEBNext Ultra DNA | Illumina HiSeq 2500 RapidMode | 150 | (Jost *et al.*, 2022) |  |
| *Hydnora esculenta* | Hesc |  | NEBNext Ultra DNA | Illumina HiSeq 2500 RapidMode | 150 | (Jost *et al.*, 2022) | SAMN44848558 |
| *Prosopanche americana* | Pame | DNeasy Plant Maxi Kit (Qiagen, Venlo, Netherlands) | Illumina TruSeq DNA | Illumina NextSeq High Output | 150 | (Jost *et al.*, 2020) | SAMN44848562 |
| *Prosopanche bonacinae* | Pbon | Modified (Doyle & Doyle, 1987) adding RNAse A (Thermo Scientific, Waltham, MA, USA) treatment (10 mg/ml) | NEBNext Ultra DNA | Illumina HiSeq RapidMode | 150 | (Jost *et al.*, 2022) | SAMN44848563 |
| *Prosopanche panguanensis* | Ppan |  | NEBNext Ultra DNA | Illumina HiSeq RapidMode | 150 | (Jost *et al.*, 2022) | SAMN44848564 |

### Table S2 Reconstruction of repetitive elements within the *H. visseri* genome

| SC* | Pipeline (RE2) provided Annotation | Annotated protein domain | Manual Annotation | SC | Pipeline (RE2) provided Annotation | Annotated protein domain | Manual Annotation |
| --- | --- | --- | --- | --- | --- | --- | --- |
| 1 | Tekay | *gag-pol* polyprotein** | *Hydnora*Tekay1 | 17 | Ogre | INT | *Hydnora*Ogre3 |
| 2 | Tekay | *gag-pol* polyprotein** | *Hydnora*Tekay2 | 18 | all | - | Ogre (unassigned) |
| 3 | Ogre | *gag-pol* polyprotein*** | *Hydnora*Ogre1 | 19 | all | GAG | *Hydnora*Ogre5 |
| 4 | all | - | *Hydnora*Ogre1 | 20 | all | - | *Hydnora*Ogre2 |
| 5 | CRM | RT | *Hydnora*CRM1 | 21 | satellite DNA | INT | *Hydnora*Galadriel1 |
| 6 | 45S rDNA | - | 45S rDNA | 22 | SIRE | RT | *Hydnora*SIRE1 |
| 7 | Angela | RH | *Hydnora*Angela1 | 23 | Ogre | GAG | *Hydnora*Ogre2 |
| 8 | CRM | INT | *Hydnora*CRM2 | 24 | Ogre | INT, RT | *Hydnora*Ogre4 |
| 9 | Tekay | RT | *Hydnora*Tekay3 | 25 | all | - | all |
| 10 | all | - | *Hydnora*Ogre2 | 26 | all | GAG | *Hydnora*Tekay4 |
| 11 | all | - | all | 27 | Ogre | INT | *Hydnora*Ogre5 |
| 12 | LTR retrotransposon | - | *Hydnora*Ogre2 | 28 | all | - | *Hydnora*Ogre3 |
| 13 | all | - | *Hydnora*Ogre1 | 29 | Tekay | INT | *Hydnora*Tekay5 |
| 14 | all | - | *Hydnora*Ogre5 | 30 | all | - | *Hydnora*Ogre2 |
| 15 | Tat | RT | *Hydnora*Ogre2 | 31 | Ogre | GAG | *Hydnora*Ogre4 |
| 16 | all | - | all | 32 | Angela | RT | *Hydnora*Angela2 |

*From the RE2 individual analysis results of *H. visseri*, superclusters comprising more than 10,000 reads (0.2% of the analyzed reads) are included in the analysis from the RE2 individual analysis result

**Showing similarity to the chromodomain (CHD), the group-specific antigen (GAG), the integrase (INT), the protease (PROT), the ribonuclease H1 (RH), and the reverse transcriptase (RT). For the principals of annotations embedded in RepeatExplorer2 (RE2), see (Novák *et al.*, 2013, 2020).

*** showing similarities to GAG, INT, PROT, RH, RT, and the archeal ribonuclease H1 (aRH)

**Table S3** Hydnoraceae short read mapping to 5S rDNA references

| Species | Number of Illumina reads | Num. of mapped reads | [%] |
| --- | --- | --- | --- |
| *H. visseri* | 95,483,059 | 291 | 0.00030 |
| *H. longicollis* | 257,184,941 | 122 | 0.00005 |
| *H. triceps* | 8,543,515 | 231 | 0.00270 |
| *H. africana* | 13,858,819 | 87 | 0.00063 |
| *H. hanningtonii* | 16,130,357 | 25 | 0.00016 |
| *H. abyssinica* | 23,408,887 | 45 | 0.00019 |
| *H. solmsiana* | 13,096,372 | 152 | 0.00116 |
| *H. esculenta* | 18,977,608 | 39 | 0.00021 |
| *P. americana*** | 186,620,441 | 265,644 | 0.14234 |
| *Aristolochia fimbriata* | 60,754,989 | 45,193 | 0.07439 |

*Proportion of reads from each *Hydnora* short read dataset that mapped to the 5S rDNA reference from *H. triceps* using Bowtie2. The reads are concordantly or disconcordantly aligned onto the paired-end reads, using mixed mode (default setting in Bowtie2). The number of aligned reads were viewed and counted using Geneious v.6.1.8. For comparison, short reads of a close photoautotrophic relative of the Hydnoraceae, *Aristolochia fimbriata* (Aristolochiaceae, SRR13748080) were mapped to the 5S rDNA reference from *H. triceps* as well.

**Table S4** NCBI identifiers for the input genome sequencing data and the reconstructed 5S rDNAs for 5S rDNA sequence comparison between Hydnoraceae and angiosperms (Separated Excel file is provided)

**Table S5** Summary of genomic proportions of repetitive elements in *Hydnora* genomes

| Repeat proportion (%) | | *H. visseri* | *H. longicollis* | *H. triceps* | *H. africana* | *H. hanningtonii* | *H. abyssinica* | *H. solmsiana* | *H. esculenta* | Mean |
| --- | --- | --- | --- | --- | --- | --- | --- | --- | --- | --- |
| Class I | Ty3-*gypsy* | 37.4 | 31.6 | 21.6 | 40.2 | 22.5 | 22.3 | 28.5 | 9.7 | 26.7 |
|  | Ty1-*copia* | 2.4 | 2.5 | 0.7 | 1.9 | 1.5 | 1.4 | 1.9 | 1.3 | 1.70 |
|  | LINEs | 0.25 | 0.23 | 0.19 | 0.07 | 0.00 | 0.25 | 0.25 | 0.00 | 0.22 |
|  | Pararetrovirus | 0.14 | 0.02 | 0.07 | 0.14 | 0.00 | 0.00 | 0.00 | 0.00 | 0.08 |
| Class II | DNA transposons | 0.00 | 0.00 | 0.00 | 0.00 | 0.10 | 0.10 | 0.23 | 0.47 | 0.23 |
| Tandem repeats | 45S rDNA | 1.41 | 1.98 | 0.54 | 0.38 | 0.07 | 0.14 | 0.12 | 0.07 | 0.59 |
|  | 5S rDNA | 0.00 | 0.00 | 0.01 | 0.00 | 0.00 | 0.00 | 0.00 | 0.00 | 0.00 |
|  | satellite DNAs | 0.00 | 0.00 | 0.05 | 0.00 | 0.47 | 0.00 | 0.02 | 0.07 | 0.13 |
| Unclassified repeats | | 14.69 | 20.38 | 26.63 | 10.7 | 14.66 | 21.9 | 18.77 | 23.38 | 18.89 |
| Total repeat proportion | | 56.25 | 56.73 | 49.77 | 52.99 | 39.34 | 46.06 | 49.78 | 34.55 | 48.18 |

**Table S6** Summary of genomic proportions of repetitive elements in *Prosopanche* genomes

| Repeat proportion (%) | | *P. americana* | *P. panguanensis* | *P. bonacinae* | Mean |
| --- | --- | --- | --- | --- | --- |
| Class I | Ty3-*gypsy* | 16.66 | 7.63 | 9.38 | 11.22 |
|  | Ty1-*copia* | 1.11 | 0.00 | 0.84 | 0.98 |
|  | Pararetrovirus | 0.00 | 0.00 | 0.00 | 0.00 |
|  | LINEs | 0.33 | 0.05 | 0.06 | 0.15 |
| Class II | DNA transposons | 2.62 | 1.84 | 7.96 | 4.14 |
|  | Helitron | 0.03 | 0.00 | 0.00 | 0.03 |
|  | 45S rDNA | 0.00 | 0.09 | 0.12 | 0.11 |
|  | 5S rDNA | 0.55 | 15.71 | 0.41 | 5.56 |
|  | satellite DNAs | 0.2 | 6.89 | 10.88 | 5.99 |
|  | Unclassified repeats | 21.57 | 14.31 | 22.82 | 19.57 |
| Total repeat proportion | | 43.07 | 46.52 | 52.48 | 47.36 |

**Table S7** Relative genomic abundance of abundant repeats in Hydnoraceae genomes

| Relative abundance of shared repeats [%] | T1 | T2 | T3 | T4 | T5 | O1 | O2 | O3 | O4 | O5 |
| --- | --- | --- | --- | --- | --- | --- | --- | --- | --- | --- |
| *H. visseri* | 17.0 | 13.3 | 18.1 | 30.0 | 13.3 | 17.0 | 22.6 | 12.9 | 22.8 | 22.2 |
| *H. longicollis* | 14.9 | 11.4 | 19.3 | 18.5 | 14.1 | 14.7 | 18.8 | 9.1 | 21.4 | 18.9 |
| *H. triceps* | 21.3 | 9.0 | 10.2 | 24.8 | 7.6 | 18.3 | 28.6 | 14.3 | 24.2 | 36.4 |
| *H. africana* | 18.6 | 10.7 | 11.7 | 26.6 | 4.1 | 21.6 | 21.8 | 16.0 | 31.5 | 22.2 |
| *H. hanningtonii* | 8.7 | 8.0 | 8.2 | 0.0 | 11.2 | 8.9 | 3.3 | 10.2 | 0.0 | 0.1 |
| *H. abyssinica* | 8.1 | 7.2 | 7.6 | 0.0 | 11.5 | 8.9 | 3.3 | 10.4 | 0.0 | 0.2 |
| *H. solmsiana* | 9.3 | 8.4 | 10.3 | 0.1 | 38.0 | 8.0 | 1.5 | 22.5 | 0.0 | 0.1 |
| *H. esculenta* | 1.5 | 5.4 | 3.1 | 0.0 | 0.0 | 2.3 | 0.0 | 0.2 | 0.0 | 0.0 |
| *P. americana* | 0.3 | 14.9 | 4.8 | 0.0 | 0.0 | 0.0 | 0.0 | 0.1 | 0.0 | 0.0 |
| *P. panguanensis* | 0.1 | 8.8 | 2.8 | 0.0 | 0.0 | 0.1 | 0.0 | 0.7 | 0.0 | 0.0 |
| *P. bonacinae* | 0.3 | 3.0 | 3.9 | 0.0 | 0.3 | 0.1 | 0.0 | 3.7 | 0.0 | 0.0 |
| *A. fimbriata* | 0.0 | 0.0 | 0.0 | 0.0 | 0.0 | 0.0 | 0.0 | 0.0 | 0.0 | 0.0 |
| Num. of reads | 286663 | 212563 | 23375 | 3200 | 5890 | 176928 | 24801 | 16044 | 11204 | 29876 |
| Relative abundance of shared repeats [%] | En/Spm_CACTA | hAT | satellite DNAs | 5S | 45S |  |  |  |  |  |
| *H. visseri* | 0.0 | 0.0 | 0.0 | 0.0 | 20.4 |  |  |  |  |  |
| *H. longicollis* | 0.0 | 0.0 | 0.0 | 0.0 | 27.9 |  |  |  |  |  |
| *H. triceps* | 0.0 | 0.0 | 0.0 | 0.0 | 13.4 |  |  |  |  |  |
| *H. africana* | 0.0 | 0.0 | 0.0 | 0.0 | 4.3 |  |  |  |  |  |
| *H. hanningtonii* | 0.0 | 0.0 | 0.0 | 0.0 | 0.8 |  |  |  |  |  |
| *H. abyssinica* | 0.0 | 0.0 | 0.0 | 0.0 | 1.9 |  |  |  |  |  |
| *H. solmsiana* | 0.0 | 0.0 | 0.0 | 0.0 | 1.3 |  |  |  |  |  |
| *H. esculenta* | 0.0 | 0.0 | 0.0 | 0.0 | 0.9 |  |  |  |  |  |
| *P. americana* | 10.5 | 9.0 | 0.0 | 3.5 | 0.2 |  |  |  |  |  |
| *P. panguanensis* | 16.9 | 70.5 | 27.1 | 94.1 | 2.0 |  |  |  |  |  |
| *P. bonacinae* | 72.6 | 20.4 | 45.0 | 2.4 | 1.5 |  |  |  |  |  |
| *A. fimbriata* | 0.0 | 0.0 | 28.0 | 0.1 | 25.4 |  |  |  |  |  |
| Num. of reads | 35514 | 6840 | 99229 | 71275 | 28658 |  |  |  |  |  |

*During the RE2 comparative analysis, the pre-labelled reads from all Hydnoraceae species are jointly clustered and annotated according to the reference repeat database (reconstructed from the *H. visseri* genome). Subsequently, the pooled reads per each repeat (‘Num. of reads’ in the table) were again classified according to the species-specific code, aiming to reveal the relative genomic abundance of shared repeats and other abundant repeats in Hydnoraceae genomes.
